# Metabolomic Investigations Reveal Lignin Deconstruction by Human Gut Microbiota

**DOI:** 10.64898/2026.10.16.759694

**Authors:** Christopher A. Dicksion, Elise A. Smentek, Daniel J. Yelle, Jonathan D. Rickmeyer, Elizabeth N. Bess

## Abstract

Lignin is a polyphenolic dietary fiber whose role in human nutrition has been debated for over a century. Herein, we reveal the ability of human gut microbiota to deconstruct lignin in anaerobic conditions. Quantitative ^13^C-tracing mass spectrometric analyses showed that sustained microbial deconstruction of lignin feeds into a pathway for the biosynthetic production of enterolignans. We used 2D NMR to characterize the microbe-induced structural changes in the lignin polymer. Untargeted mass spectrometry demonstrated that human gut microbiota not only deconstruct lignin but also that they do so on rapid timescales. While most existing reports of anaerobic lignin metabolism have solely focused on the production of small metabolites, we additionally profiled the deconstruction of large lignin oligomers. Moreover, we identified the putative structure of a large lignin oligomer and traced its metabolism to downstream metabolites. Collectively, we provide a compelling body of evidence to support the hypothesis that human gut microbiota deconstruct lignin and to establish the human gut microbiome in the paradigm of anaerobic lignin deconstruction.

## Introduction

Lignin is a dietary fiber consumed in multi-gram quantities in minimally processed plant-rich diets,^1^ yet its fate in the human digestive tract has largely remained a mystery.^2,3^ A polymer of polyphenols, lignin is a primary structural component of plant cell walls that accounts for ∼30% of all organic carbon in the biosphere.^4^ Plants assemble lignin via radical coupling of *para*-coumaryl alcohol, coniferyl alcohol, and sinapyl alcohol, forming irregular, heterogenous polymers with carbon–carbon and aryl–ether couplings. Although lignin’s chemical composition gives rise to its recalcitrance to biotic degradation,^5^ investigations of soil microbes show that an astonishing array of aromatic metabolites with bioactive properties can result from lignin’s microbial degradation.^6,7,8^ We sought to reveal if anaerobic human gut bacteria can also deconstruct lignin and what metabolites may result from this process.

In environmental settings, aerobic degradation of lignin is well characterized, particularly with respect to white-rot fungi.^9^ These organisms employ a suite of lignin-degrading enzymes, including peroxidases and laccases.^10^ The enzymatic mechanisms are strictly oxygen dependent, relying on the generation of highly reactive oxygen species or the reduction of molecular oxygen to break the stable carbon–carbon and aryl–ether bonds of the lignin polymer.^11^ This context has given rise to the prevailing hypothesis that lignin deconstruction mediated by human gut microbiota is not possible because the oxygen-dependent ligninolytic chemistry found in aerobic environments is unlikely in the largely anaerobic environment of the human gut.^12,13^

Challenging the notion that lignin is entirely recalcitrant under anaerobic conditions, emerging research is providing evidence of anaerobic lignin degradation. Li et al. demonstrated that a species of fungus-cultivating termite deconstructs lignin polymer with a whole-gut transit time under 4 hours.^14^ Begum et al. utilized synthetic, deuterated lignin polymers to trace their metabolic fate in rats, showing that lignin can be metabolized in the anaerobic environment of the murine gut.^15^ Lankiewicz et al. identified anaerobic gut fungi common in herbivores that are capable of deconstructing lignin.^16^ Kasimir et al. anaerobically cultured microbiota derived from pig ceca to show that ^13^C-labelled lignin was degraded into a variety of phenolic acids, lignans, and enterolignans within 72 hours.^17^ Altogether, recent evidence coheres to establish a new paradigm: anaerobic gut environments contain microbiota that not only deconstruct lignin to produce bioactive metabolites but do so on rapid timescales.

Whether human gut microbiota deconstruct lignin has remained an outstanding question. This complex anaerobic microbial community is sampled via fecal material, which contains lignin due to its ubiquitous presence in minimally processed dietary plants. Prior investigations have been hampered by the confounding effects of lignin and its metabolites that are endogenous to the fecal material.^18,19^ We hypothesized that we could circumvent this challenge and definitively trace metabolism of lignin by human gut microbiota using uniformly ^13^C-labelled (per-^13^C) lignin, as any products of lignin deconstruction would be labeled with this rare isotope. Here, we report our use of per-^13^C lignin to quantify the flux of metabolites in a lignin-related biosynthetic pathway, characterize microbe-induced structural changes to the lignin polymer by 2D NMR, reveal the metabolomic signatures of lignin deconstruction, and elucidate putative molecular structures in a metabolic pathway that leads from large lignin oligomers to monomeric phenolics.

## Materials and Methods

### Chemicals

(–)-Enterolactone, (+)-lariciresinol, (+)-pinoresinol, and (–)-secoisolariciresinol were purchased from Sigma Aldrich in 95% purity. Internal standards (IS) ^13^C_3_-rac-enterolactone and *d*_6_-rac-secoisolariciresinol were purchased from Toronto Research Chemicals in 95% purity. DMSO-*d*_6_ was purchased from Sigma Aldrich with 99.96% atom D. L-cysteine HCl (95%), HPLC grade methyl *tert*-butyl ether (MtBE), LC–MS grade acetonitrile, and LC–MS grade acetic acid were purchased from Fisher Scientific. Sterile DMSO was purchased from ATCC. Gifu anaerobic medium (GAM) was purchased from HiMedia Labs. M9 salt powder was purchased from Sigma Aldrich. Organosolv wheat lignin was obtained as a gift from the Fraunhofer Center for Chemical-Biotechnological Processes (Leuna, Germany). Natural isotope abundance (^12^C) and uniformly-^13^C labelled lignin (per-^13^C) were obtained from IsoLife. Double deionized water (DDI H_2_O) was obtained from an Elga PURELAB flex water purifier at greater than 18.2 MΩ⋅cm.

### Preparation of Cultures for Anaerobic Incubation

Fresh fecal samples were obtained from 5 healthy human volunteers. Each participant consented to participate. The study was approved by the University of California, Irvine Institutional Review Board (protocol # 20195091). Participants collected the samples with GutAlive anaerobic microbiome collection kits (Microviable, Gijón, Spain), capable of maintaining samples in an anaerobic state until they were delivered to our laboratory within 18 hours of collection. After delivery to the laboratory, the samples were transferred to an anaerobic chamber (Coy Laboratory Products, Grass Lake, MI) containing an atmosphere of 20% CO_2_, 2%–5% H_2_, and the remainder N_2_. Samples were mixed with a spatula to homogenize and then added to 15 mL conical tubes as 0.5–2.0 g aliquots before being stored at -80 °C.

For use, frozen fecal samples from 5 human donors were brought into an anaerobic chamber to thaw at room temperature for 30 minutes. The samples remained capped and were weighed in an aerobic environment before being returned to the anaerobic chamber, where 0.5 g from each of the 5 donors were mixed together in 15 mL conical tubes, in triplicate. Samples were resuspended in 12.5 mL of freshly prepared and equilibrated M9 media supplemented with 0.1% (w/v) filter-sterilized L-cysteine HCl to make a 20% (w/v) fecal suspension. The suspension was gently shaken for 5 minutes at 37 °C and then allowed to settle for 5 minutes. An aliquot of 1.4 mL of the fecal suspension was used to inoculate freshly prepared and equilibrated Gifu anaerobic media (GAM), yielding a 18.9% (v/v) suspension in the culture. An appropriate volume of M9 salts supplemented with 0.1% (w/v) L-cysteine HCl was used for sterile control conditions.

Lignin stocks were prepared in sterile DMSO at 50 mg/mL. Initial studies were performed with organosolv wheat lignin (provided by the Fraunhofer Center for Chemical-Biotechnological Processes; Leuna, Germany). Stable-isotope labelling experiments combined equal amounts of natural isotope abundance lignin and per-^13^C-lignin (IsoLife N-40420 and U-40420, respectively), except for the cultures prepared for NMR, where only per-^13^C-lignin was used. A 74 µL aliquot of the lignin stock was used to prepare 0.05% (w/v) lignin in each culture. An equivalent volume of sterile DMSO was used for control samples.

Four culture conditions were devised: GAM containing fecal microbiota and lignin (GAM_+fecal/+lignin_), GAM containing fecal microbiota but without lignin (GAM_+fecal/-lignin_), GAM without fecal microbiota but with lignin (GAM_-fecal/+lignin_), and GAM without either fecal microbiota or lignin (GAM_-fecal/-lignin_). Cultures were prepared in triplicate as 7.4 mL mixtures in Hüngate tubes that were tightly capped with butyl rubber stoppers and screw caps. Cultures were incubated at 37 °C in an anaerobic chamber with gentle shaking. Aliquots were taken at each timepoint by inverting each tube twice before drawing 600 µL of the mixture with a 22 G needle affixed to a 1 mL syringe. Each aliquot was dispensed into a tightly capped 2 mL glass HPLC vial. Samples were immediately removed from the anaerobic chamber and heated at 100 °C for 15 minutes before being chilled on ice for 15 minutes. Samples were then further aliquotted at 300 µL into 2 mL glass HPLC vials and capped before being stored at -80 °C until further processing.

Sample preparation for mass spectrometric analysis was based on an existing method.^20^ Briefly, a 300 µL aliquot of each sample was thawed at room temperature for 15 minutes before the addition of 300 µL of double deionized (DDI) H_2_O containing internal standards (^13^C_3_-rac-enterolactone and *d*_6_-rac-secoisolariciresinol, 20 nM each). Liquid–liquid extraction was performed by adding 1.2 mL MtBE-containing 0.1% (v/v) acetic acid. The samples were mixed at 1000 rpm for 5 minutes, then centrifuged at 3000 rpm for 10 minutes. A 900 µL aliquot of the MtBE layer was transferred to a glass culture tube and evaporated under N_2_ at 40 °C for 15 minutes. The residues were resuspended in 450 µL DDI H_2_O containing 20% acetonitrile, capped with a butyl rubber stopper, and mixed at 500 rpm for 15 minutes. The samples were used immediately for quantitative LC–MS/MS analysis. Upon completion of the LC–MS/MS analysis, the samples were stored at -80 °C until the execution of LC–QTof–MS analysis.

### 2D HSQC NMR Analysis

Cultures for NMR analysis were prepared in an anaerobic chamber by resuspending 0.5 g of fecal material from each of 4 donors in 10 mL of freshly prepared and atmosphere-equilibrated M9 media supplemented with 0.1% (w/v) filter-sterilized L-cysteine HCl to make a 20% (w/v) fecal suspension. The suspension was gently shaken for 5 minutes at 37 °C and then allowed to settle for 5 minutes. An aliquot of 1.4 mL of the fecal suspension was used to inoculate freshly prepared and equilibrated Gifu anaerobic media (GAM), yielding 7.4 mL of a 18.9% (v/v) suspension, including 74 µL of a 50 mg/mL stock of per-^13^C lignin in sterile DMSO (0.05 % w/v). After 96 hours of incubation, the samples were removed from the anaerobic chamber and heated at 100 °C for 15 minutes before being placed on ice for 15 minutes. To separate the insoluble components in the culture from the soluble media, a 1 mL aliquot of each culture was added to its own microcentrifuge tube and centrifuged at 15,000 rpm for 10 minutes. The supernatants were decanted into clean, 15 mL conical tubes. Another 1 mL aliquot was then added into the same microcentrifuge tube and centrifuged again. This process was repeated until the entire culture volume had been processed, at which time the pellets were stored at -80 °C for 24 hours. Frozen pellets were lyophilized for 96 hours.

Each lyophilized pellet was weighed separately (GAM_-fecal/+lignin_ = 8.9 mg, GAM_+fecal/+lignin_ = 8.4 mg) and then dissolved directly into its own Shigemi NMR tube (susceptibility matched plunger to DMSO-*d_6_*) using 200 µL of DMSO-*d_6_* and sonicated at 35 °C for 1 hour. The dissolved samples were homogenous gel solutions. NMR spectra were acquired on a Bruker-Biospin (Rheinstetten, Germany) AVANCE III HD 500 MHz spectrometer fitted with a nitrogen-cooled 5 mm Prodigy TCI gradient cryoprobe with inverse geometry. One-bond ^1^H–^13^C correlation (HSQC) spectra were acquired using adiabatic Bruker pulse program hsqcetgpsisp2.2 and processed as previously described^21^ using Bruker TopSpin 4.5.0 software.

### Quantitative Analysis by LC–MS/MS

Quantitative mass spectrometry (LC–MS/MS) was performed with a Waters Xevo TQ Absolute mass spectrometer coupled to a Waters Acquity I-Class Premier UPLC system (Waters, Milford, MA, USA). The instrument was operated in multiple-reaction-monitoring and negative-electrospray-ionization modes according to precursor, fragment, and voltage parameters that were previously established (**Table S1**).^20^ Per-^13^C fragment *m/z* values were established by using CFM-ID (V. 4.0)^22^ to identify existing ^12^C fragment structures and then calculating the monoisotopic per-^13^C *m/z* such that all carbon atoms in the fragment were ^13^C isotopes.^23^ A 2 µL injection was separated with a Waters Acquity Premier BEH C18 column (2.1 x 50 mm, 1.7 µm particle size) equipped with a VanGuard FIT BEH C18 guard column. The column was maintained at 50 °C. Solvent A was DDI H_2_O with 0.1% (v/v) acetic acid, and Solvent B was acetonitrile with 0.1% (v/v) acetic acid running at a flow rate of 0.500 mL/min. The solvent gradient was 20% B to 40% B from 0–3.5 minutes, then ramped to 95% B over 0.1 minutes and held isocratic at 95% B from 3.6–4.15 minutes, and finally reduced to 20% B from 4.15–5.0 minutes.

### Analysis of Unknown Metabolites via LC–MS

Untargeted mass spectrometry (LC–QToF-MS) was performed with a Waters Xevo G2-XS mass spectrometer coupled to a Waters Acquity LC system. A 10 µL aliquot was injected onto an Acquity Premier CSH Hexyl-Phenyl column (2.1 x 100 mm, 1.7 µm particle size) heated to 50 °C. Metabolites were eluted with DDI H_2_O containing 0.1% (v/v) acetic acid (Solvent A) and acetonitrile containing 0.1% (v/v) acetic acid (Solvent B) running at a flow rate of 0.400 mL/minute according to the following gradient: 100% A from 0–1 minute, a ramp to 10% B from 1–2 minutes, main elution via a ramp to 60% B from 2–13 minutes, washout with a ramp to 100% B from 13–15 minutes and then held constant until 17 minutes, followed by re-equilibration at 100% A from 18–20 minutes. The capillary voltage was 2.5 kV, and the sampling cone was set to 40 V. The source temperature was 100 °C, with desolvation gas flow of 800 L/Hr at 350 °C. Centroid mass spectra were collected using the fast data dependent acquisition (FDDA) and ESI^−^ modes, with 1 precursor scan every 0.2 seconds across a 50–1000 Da mass range, and 4 DDA fragment scans for 0.2 seconds each for selected precursors. A voltage ramp from 10–100 V was applied for collision across the entire mass range. Leucine enkephalin (100 pg/µL) was sprayed into the source at 30 second intervals as an external lock-mass.

Additional fragmentation spectra were obtained for some features via targeted fragmentation analysis using a Waters Xevo TQ Absolute mass spectrometer. Briefly, methods were configured to target *m/z* values for particular ^12^C/^13^C pairs that had been identified during the metabolic network analysis. The cone voltage was 10 V, and the collision energy was ramped from 10 eV to 40 eV across the *m/z* range 50–600 Da. A 10 µL bolus was injected with a Waters Acquity I-Class Premier UPLC system and separated with a Waters Acquity Premier BEH C18 column (50 x 2.1 mm, 1.7 µm particle size) equipped with a VanGuard FIT BEH C18 guard column. The column was maintained at 50 °C. Solvent A was DDI H_2_O with 0.1% (v/v) acetic acid, and Solvent B was acetonitrile with 0.1% (v/v) acetic acid running at a flow rate of 0.500 mL/min. The solvent gradient was 20% B to 40% B from 0–3.5 minutes, then ramped to 95% B over 0.1 minutes and held isocratic at 95% B from 3.6–4.15 minutes, and finally reduced to 20% B from 4.15–5.0 minutes.

### Processing and Analysis of Mass Spectrometric Data

The TargetLynx module of MassLynx (V. 4.1) was used to quantify LC–MS/MS data. MS-Dial (V.5.5.251021) was used to perform peak alignment and feature assignment (*m/z* and retention time) for each of the samples. The parameters included the default settings, except for MS^1^ tolerance of 0.01 Da and MS^2^ tolerance of 0.025 Da, minimum peak height of 1000 with a slice width of 0.1 Da, and default values for spectrum deconvolution. Feature alignment was performed against a mixed sample (5 µL from each of the GAM_+fecal/+lignin_ and GAM_+fecal/-lignin_ samples), with retention time tolerance of 0.02 minutes and MS^1^ tolerance of 0.02 Da. Normalization of each feature was performed by subtracting the mean intensity of the instrument blanks and the GAM_-fecal/-lignin_ controls, and 0 or negative values were imputed with 10 (a value more than 5 times lower than the lower limit of detection) to avoid computational issues that arise from missing or zero values in quotient or log mathematical transformations.^24^

Manual curation of the mass spectra was performed to identify pairs of ^12^C and ^13^C metabolites with the same molecular formulas. Individual MS^1^ scans were sequentially inspected across all replicates and timepoints of the GAM_+fecal/+lignin_ samples to identify the *m/z* and retention times of each feature in a ^12^C/^13^C pair. Molecular formulas for ^12^C/^13^C feature-pairs were assigned by first identifying the number of carbon atoms in the metabolite as the nominal difference between the ^13^C monoisotopic *m/z* and the ^12^C monoisotopic *m/z*. Then, the observed monoisotopic *m/z* for each ^12^C feature was entered into the monoisotopic mass module of the web-based chemcalc platform (chemcalc.org) with default settings except for the allowance of up to 50 oxygen atoms.^23^ The results were filtered according to the number of carbon atoms, and the molecular formula with both the correct number of carbon atoms and the smallest deviation from the theoretical value was selected. The molecular formula for each feature was then modified to contain all ^13^C atoms before being input into the molecular formula module of the chemcalc platform to obtain the theoretical *m/z* for each per-^13^C feature.

Kendrick mass defect analysis was performed according to **Equations 1–2** to assess the ^12^C/^13^C pairs, using 14.01565 Da as the monoisotopic *m/z* for a ^12^CH_2_ subunit and 15.01900 Da as the monoisotopic *m/z* for a ^13^CH_2_ subunit.

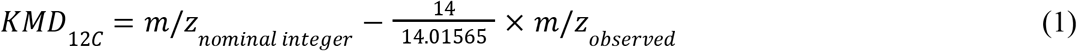

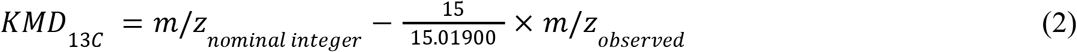

Van Krevelen analysis was performed by comparing the H/C and O/C atom ratios from the molecular formulas for each feature to the corresponding H/C and O/C ranges established by Laszakovits and MacKay^25^ or by Oesterle et al.^26^

MetaboAnalyst (V. 6.0)^27^ was used to perform principal component analysis and hierarchical clustering analysis (heatmap). Briefly, per-^13^C feature lists were uploaded for single factor statistical analyses. The mass tolerance was set to 0.01 Da, and the retention time tolerance was set to 0.05 minutes. Data filtering was configured such that no features were filtered out. The data were transformed with log_10_ transformation and Pareto scaling. The heatmap was generated according to the *z*-score for each replicate of each feature across all conditions, according to **Equation 3**.

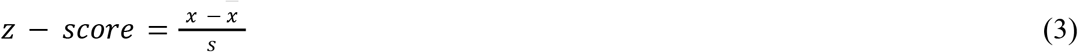

In this equation, *x* is the observed intensity in a given replicate, *x̄* is the mean intensity across all replicates (n = 3), timepoints, and conditions, and *s* is the standard deviation. GraphPad Prism (V. 11.0) was used to perform a simple linear regression for Kendrick mass defect analysis (default settings), to perform area under the curve (AUC) calculations for each replicate of each per-^13^C feature in each condition across the entire 96 hour incubation (baseline of *y* = 0, no ignored peaks, and the requirement that all peaks be above the baseline), and to perform multiple unpaired *t*-tests on various data sets (parameters: assume a normal distribution of samples from populations with equal standard deviations and statistical discoveries were pursued according to a false discovery rate approach based on the two-stage step-up method of Benjamini, Krieger, and Yekutieli multiple-testing correction (false discovery rate Q=1%). Net area under the curve calculations were performed for each per-^13^C feature according to **Equation 4**, such that the mean AUC for the GAM_-fecal/+lignin_ control condition was subtracted from the mean AUC for the GAM_+fecal/+lignin_ condition (n = 3).

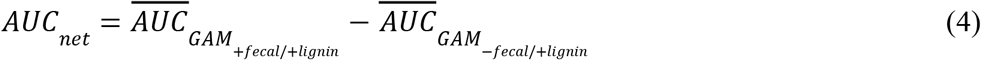

## Results and Discussion

### Isotope-Tracing Tracks Metabolites of Human Gut Bacterial Lignin Deconstruction

Lignin’s heterogeneous composition of monolignols is a challenge to analytical assessment of its biochemical deconstruction (**Figure 1A**). Without knowledge of how the polymer is digested nor the metabolites that result, a quantifiable signal of its deconstruction has remained elusive. To overcome this, we recognized that the characteristic β–β monolignol inter-unit bond comprises 2–5% of monolignol inter-unit bonds found in wheat straw.^28^ This β–β linkage is prominent in pinoresinol, a motif embedded in the lignin polymer. Once freed from the polymer, pinoresinol is a feedstock of the well-characterized gut bacterial biosynthesis of enterolactone,^29^ a molecule known for its anti-breast cancer effects.^30^ Because enterolactone is a terminal metabolite in the biochemical cascade that begins with pinoresinol, we hypothesized that accumulation of enterolactone could serve as a proxy for lignin deconstruction in vitro (**Figure 1A**).^15^ Indeed, Begum et al. demonstrated that gut microbiota native to rats degraded poly-deuterated synthetic dehydrogenation lignin polymers to di-deuterated enterolactone in vivo.^15^

**Figure 1:**
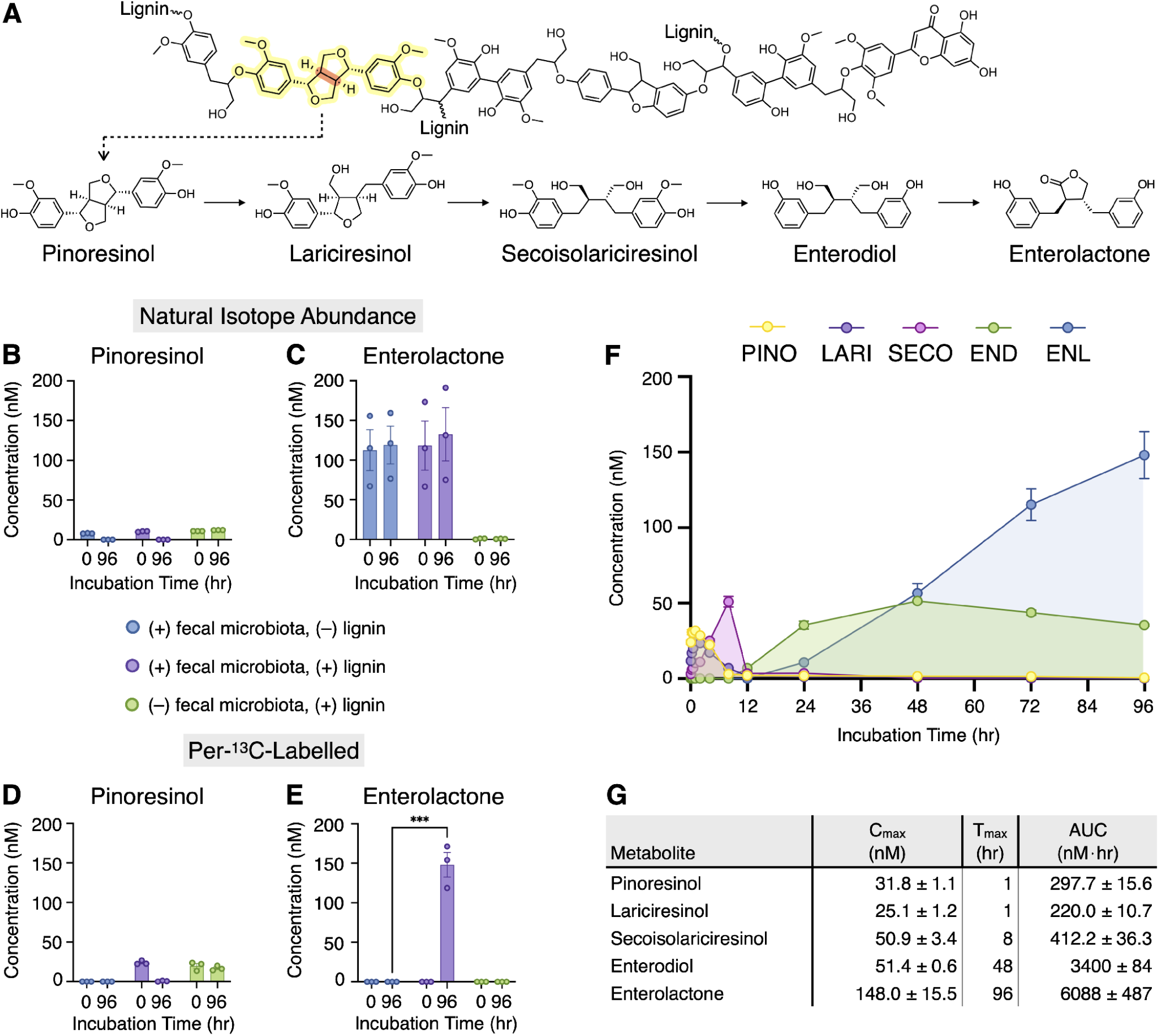
Quantitative metabolomic analyses suggest lignin deconstruction by human gut microbiota. **(A)** Chemical structures of a plausible lignin polymer and of metabolites in a lignin-associated biochemical pathway, with lignin-embedded pinoresinol highlighted in yellow and its β–β bond emphasized in orange. **(B–E)** Quantitative analysis of pinoresinol and enterolactone after 0 hours and 96 hours when human gut microbiota were anaerobically incubated with **(B–C)** natural isotopic abundance lignin or **(D–E)** per-^13^C lignin (mean ± standard error of the mean, n = 3; unpaired *t*-test, *p*-adjusted = 0.0007). Discovery was determined by multiple unpaired *t*-tests with the two-stage step-up method of Benjamini, Krieger, and Yekutieli multiple-testing correction (false discovery rate Q=1%). **(F)** Concentration profiles of per-^13^C-labelled metabolites—pinoresinol (PINO), lariciresinol (LARI), secoisolariciresinol (SECO), enterodiol (END), and enterolactone (ENL)—when human gut microbiota were incubated with per-^13^C lignin (mean ± standard error of the mean, n = 3). **(G)** Kinetic measurements of lignan and enterolignan metabolism derived from results in **(F)**. C_max_ is the maximum concentration, and T_max_ is the incubation time at which C_max_ was observed. AUC is the mean area under the curve. C_max_ and AUC are presented as the mean ± standard error of the mean. See **Tables S1–S3** for source data.

Envisioning enterolactone production as a signal of lignin deconstruction, we incubated human fecal samples with lignin. Fecal samples (5 donors) were collected using GutAlive Microviable kits that preserve bacteria in an anaerobic environment from the time of collection until receipt in our anaerobic chamber, where samples were aliquoted and then stored at -80 °C until use. Samples were incubated anaerobically for 96 hours in GAM, according to four experimental conditions: media only (negative control, GAM_-fecal/-lignin_), with lignin (0.05%, w/v) but without human gut microbiota (negative control, GAM_-fecal/+lignin_), without lignin but with human gut microbiota (negative control, GAM_+fecal/-lignin_), and with lignin (0.05%, w/v) and with human gut microbiota (GAM_+fecal/+lignin_). Pinoresinol and enterolactone were quantified using our established LC–MS/MS method at the beginning (t = 0 hours) and end (t = 96 hours) of the incubation period (**Figure 1B–C**).^20^ While no significant differences in pinoresinol or enterolactone levels were observed in bacterial cultures as a function of lignin supplementation, the substantial enterolactone present at the start of culturing (t = 0 hours) suggested that enterolactone and lignin endogenous to the human fecal samples masked enterolactone production resulting from deconstruction of supplemented lignin.

Longitudinal quantification of lignin deconstruction required differentiation between lignin endogenous to the fecal sample and lignin supplemented at culture initiation. We accomplished this using uniformly ^13^C-labelled lignin (per-^13^C-lignin). Human fecal samples were again anaerobically cultured in nutrient-rich media as a function of a 1:1 mixture of per-^13^C lignin (isolated from wheat grown in a ^13^CO_2_ atmosphere)^31^ and natural isotope abundance lignin. Per-^13^C pinoresinol and enterolactone as well as intermediates in this pathway (lariciresinol, secoisolariciresinol, and enterodiol) were longitudinally quantified (12 timepoints from 0 to 96 hours) by LC–MS/MS using our established method.^20^ At the start of incubation (t = 0 hours), per-^13^C pinoresinol was present at low concentrations (20–25 nM) and per-^13^C enterolactone was below the limit of quantitation (0.244 nM) in the cultures supplemented with lignin (**Figure 1D–E**). Following a 96-hour incubation of per-^13^C lignin with fecal microbiota, per-^13^C enterolactone reached 148.0 ± 15.5 nM. In contrast, per-^13^C enterolactone was neither detected in microbiota-free samples incubated with per-^13^C lignin nor in cultures of fecal microbiota absent per-^13^C lignin (**Figure 1E**).

Metabolic flux in the per-^13^C pinoresinol-to-enterolactone conversion suggests that lignin supplied a continuous feedstock to this system. The rapid initial concentration increase in per-^13^C pinoresinol followed by lariciresinol and then by secoisolariciresinol aligned with the reported metabolic order of this conversion (**Figure 1A**).^32^ By 12 hours, per-^13^C enterodiol concentration increased, until an approximately steady-state concentration was maintained from 24 to 96 hours. This corresponded with low steady-state concentrations of per-^13^C pinoresinol, lariciresinol, and secoisolariciresinol that hovered near the lower limit of quantitation (0.98 nM) over the same period. Despite the concentrations of per-^13^C pinoresinol, lariciresinol, secoisolariciresinol, and enterodiol remaining near constant from 24 to 96 hours, these molecules’ downstream product—per-^13^C enterolactone—steadily increased in concentration at a near-constant rate.

The continuous production of per-^13^C enterolactone requires a continuous supply of its metabolic precursors. The steady-state concentration of pinoresinol, lariciresinol, and secoisolariciresinol indicate that each is rapidly metabolized, until the transformation to enterodiol. The accumulation of enerodiol indicates that the final step in this pathway—conversion of enterodiol to enterolactone—is rate-determining, an observation that aligns with accumulation of enterodiol relative to enterolactone in studies of mice^32^ and human gut microbiota.^33^ Taken together, these data suggest that it is the gut bacterial deconstruction of per-^13^C lignin that affords the continuous feedstock to form per-^13^C enterolactone. These findings also motivated subsequent structural analysis of lignin (2D NMR) and comprehensive untargeted isotope-tracing metabolomics to reveal other chemical species involved in gut bacterial deconstruction of lignin.

### 2D NMR Demonstrates that Human Gut Bacteria Chemically Modify Lignin

Due to the robust production of per-^13^C enterolactone from per-^13^C lignin cultured with human fecal bacteria, we suspected that evidence of this deconstruction would be reflected in the chemical structure of the lignin polymer itself. The heterogeneous structure and molecular weight of lignin precluded its characterization through the targeted LC–MS/MS method described above; however, structural characterization of lignin via 2D NMR is well established.^28,34^

Chemical alterations made by human gut bacteria to the lignin polymer were assessed using HSQC NMR spectroscopy. Fecal samples from 4 individuals were combined and incubated in nutrient-rich media appended with per-^13^C lignin (0.05%, w/v). An additional sample was identically prepared except no fecal sample was added, serving as a sterile control. After 96 hours of anaerobic incubation, insoluble particles (including bacteria and lignin) were separated from the supernatant via centrifugation and then lyophilized. The lyophilized samples were then solubilized in DMSO-*d*_6_ at approximately 40 mg/mL, producing a viscous gel solution. Samples were prepared in Shigemi NMR tubes because they allow for the sample concentration to be increased 2–3 fold, leading to an increase in the signal-to-noise ratio when subjected to an HSQC pulse sequence.

Analysis of the 2D NMR spectra (**Figures 2A–B**) indicated that the lignin polymer is deconstructed by human gut bacteria. This was assessed by tracking the chemical motifs common to the lignin polymer, which are color-coded in **Figure 2C** to match the contours in **Figures 2A–B** that indicate the appropriate structure and position of the motifs. One such structural motif is β-aryl ether (β-O-4), which is a primary lignin linkage found in wheat straw that accounts for approximately 75% of all interunit linkages.^34^ Lignin incubated with or without bacteria shows a strong presence of β-O-4 ether structures (**Figure 2A–B**), with all three linkage positions depicted (A_ɑ_, A_β_, and A_ɣ_, **Figure 2C**). In the absence of bacteria, integration of the β-O-4 ether A_ɑ_ contour cluster, relative to the lignin aryl methoxy (OMe) contour, yielded a frequency of 0.0040 units per OMe; the analogous frequency in lignin incubated with gut bacteria was 0.0038 units per OMe. The decreased frequency of β-O-4 ether linkages detected when lignin was incubated with fecal bacteria indicates that bacteria degraded lignin at the β-O-4 ether linkages.

**Figure 2:**
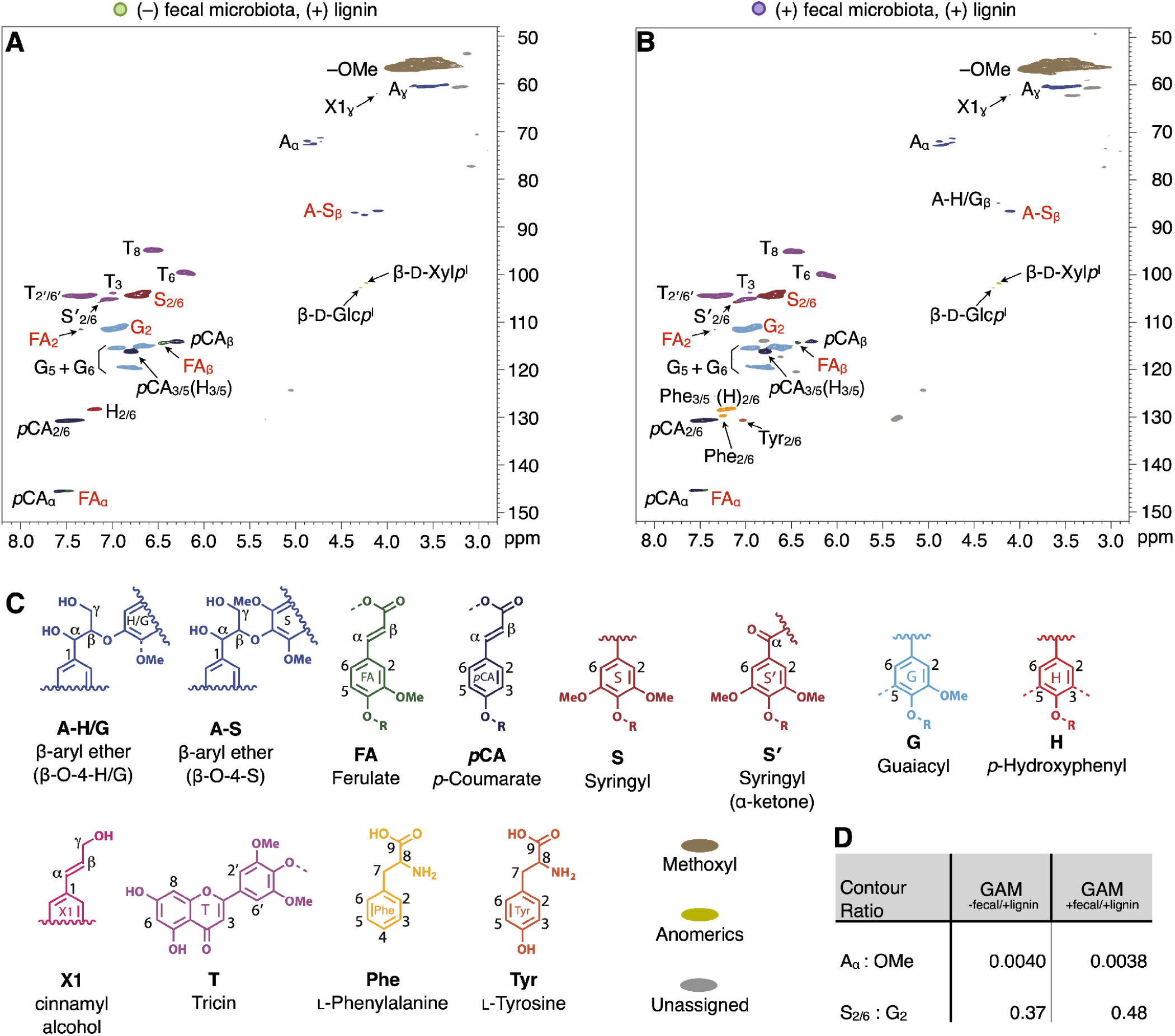
NMR analysis suggests cleavage of guaiacyl β-aryl–ether linkages in lignin polymers by human gut microbiota. One-bond ^1^H–^13^C correlation (HSQC) NMR spectra of lignin pellets after per-^13^C lignin was anaerobically incubated at 37 °C for 96 hours **(A)** without microbiota or **(B)** with microbiota in nutrient-rich media (n = 1 sample made from a fecal mixture of n = 4 donors); labels of guaiacyl and syringyl contours with differential intensity between sample conditions are highlighted in red. **(C)** Color-coded structural motifs for contours observed in the spectra. **(D)** Contour ratios between culture conditions GAM_-fecal/+lignin_ and GAM_+fecal/+lignin_ that are diagnostic for structural modifications of lignin.

Alterations to the lignin backbone, such as β-O-4 ether cleavage, are commonly assessed using the OMe contour as a reference because this contour is well-defined in the NMR spectra.^21^ Additionally, the lignin OMe is typically considered to be less chemically labile than the lignin backbone. However, the typical OMe signal stability may not exist in the presence of human gut bacteria, which demethylate aryl methoxys in small molecules.^35^ Whether these bacteria demethylate lignin polymer has remained unknown. Nonetheless, even if lignin is simultaneously undergoing demethylation and β-O-4 ether cleavage, the bacteria-driven decrease in β-O-4 ether:OMe indicates that β-O-4 ether is at least degraded faster than OMe.

We further characterized the nature of gut bacterial β-O-4 ether cleavage by quantifying the ratio of syringyl and guaiacyl units (**Figure 2C**), which are the dominant aryl motifs in wheat straw lignin.^34^ In the absence of bacteria, the syringyl:guaiacyl ratio was estimated to be 0.37 by integrating the S_2/6_ and G_2_ contours; the S_2/6_ integral was logically divided by two because each syringyl unit has two protons, compared to the single proton of a guaiacyl unit. Lignin incubated with fecal bacteria displayed a syringyl-to-guaiacyl unit ratio of 0.48. The 30% increase in syringyl:guaiacyl in the presence of bacteria indicated that lignin degradation had occurred with preferential cleavage of β-O-4 ether linkages at guaiacyl units over syringyl units, with the possibility that demethylation of guaiacyl methoxy can also impact this ratio.

Additional lignin motifs that were clearly detected in both the presence and absence of bacteria were the cinnamyl alcohol end-groups (X1) and hydroxycinnamates, such as *p*-coumarates (*p*CA) and ferulates (FA). The bacteria-free samples showed a strong *p*-hydroxyphenyl unit (H) signal, but the H_2/6_ position was obscured in the bacterial culture by the amino acid phenylalanine (Phe_3/5_). The signals from phenylalanine (Phe) and tyrosine (Tyr) were only detected in samples with bacteria, likely originating from the fecal inoculum. Tricin, a common aromatic structure found as an end-unit in lignin from herbaceous plants like wheat,^34^ shows a strong presence in samples incubated with and without bacteria. Tricin is coupled via its syringyl motif in a β-O-4 ether linkage in lignin, so cleavage of its β-O-4 ether bond represents one example where bacterial deconstruction of links to syringyl units is prominent. In support of this finding, Gall et al. found that aerobic β-ether reductase enzymes in soil bacteria cleave β-O-4 bonds between tricin and hydroxypropanovanillone.^36^ Collectively, NMR analyses revealed gross structural changes of lignin polymer mediated by human gut microbiota upon anaerobic incubation, providing motivation for in-depth metabolomic analyses with mass spectrometry to profile the scope of resultant metabolites.

### A Wide Array of Metabolites Results from Human Gut Bacterial Metabolism of Lignin

Analysis by NMR allowed us to characterize the bulk changes in the lignin polymer, and tracing the conversion of lignin to enterolactone via LC–MS/MS provided a handle to definitively assess lignin deconstruction. However, we expected that this metabolism represents only a portion of the human gut microbiota’s capacity to metabolically engage lignin. Therefore, we performed an untargeted mass spectrometric analysis on the same samples that were used to quantify the conversion of lignin to enterolactone. In short, 3 experimental conditions and 1 media-only condition were employed, each in triplicate incubations: GAM_+fecal/+lignin_, GAM_-fecal/+lignin_, GAM_+fecal/-lignin_, and GAM_-fecal/-lignin_, where fecal samples were prepared as a pooled mixture of n = 5 donors and lignin was present as a 1:1 mixture of ^12^C and per-^13^C lignin. These cultures were incubated in anaerobic conditions, and aliquots were harvested at 12 timepoints between 0 and 96 hours. Statistical analyses of mass spectrometric features demonstrated that human gut microbiota deconstruct lignin (**Figure 3**).

**Figure 3:**
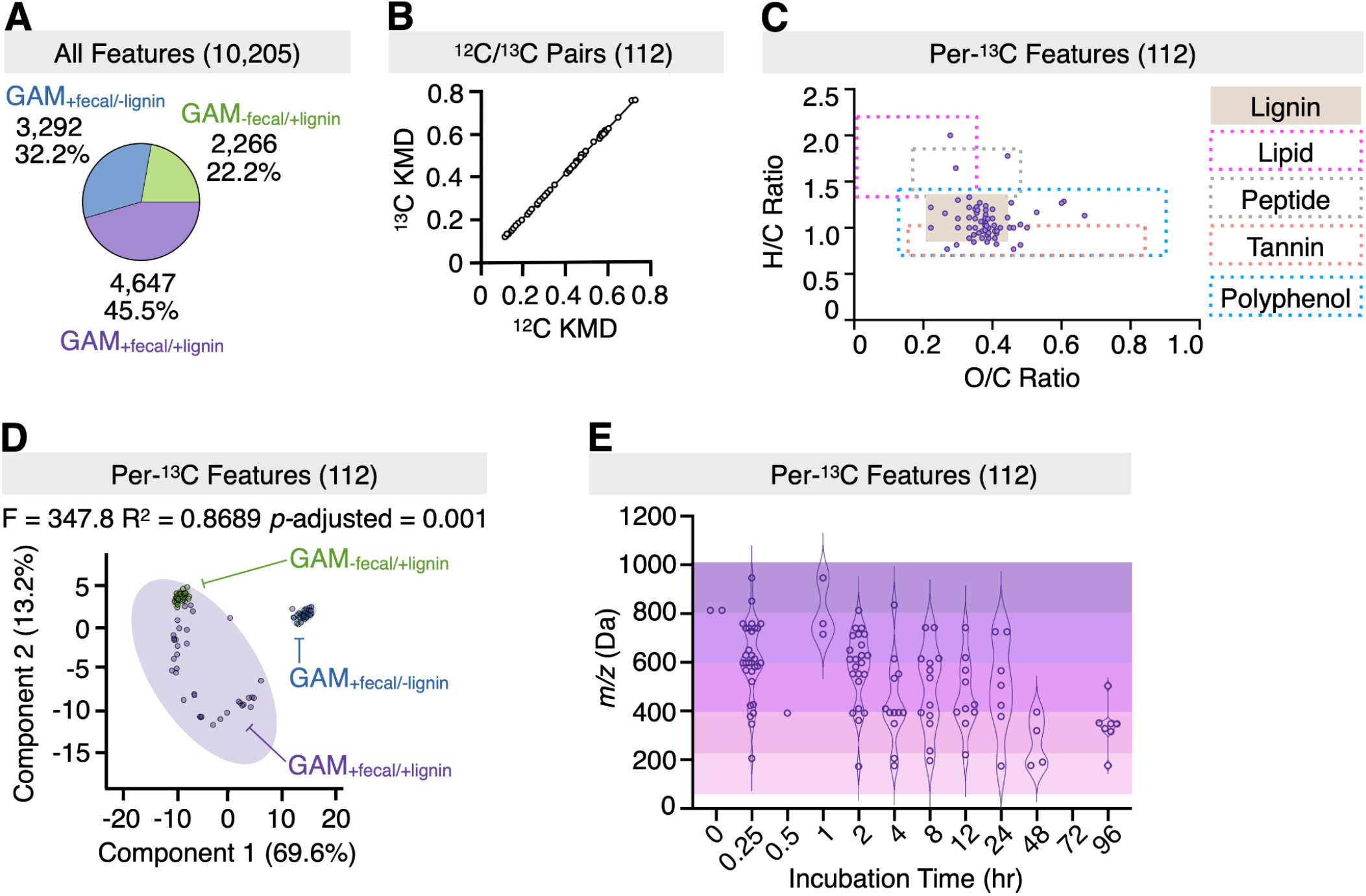
Mass spectrometric feature distribution from LC–MS analysis of cultures suggests lignin deconstruction by human gut microbiota. Cultures (n = 3) were independently prepared as fecal slurries from 5 donors and incubated anaerobically for 96 hours. (A) Distribution of mass spectrometric features identified by MS-Dial (V5.5.251021) consisting of paired retention time and *m/z* values. Features were normalized by subtracting the mean intensity from the instrument blanks and media-only control samples (GAM_-fecal/-lignin_), finding 10,205 features in the GAM_+fecal/-lignin_, GAM_+fecal/+lignin_, and GAM_-fecal/+lignin_ conditions. From the mean signal intensity for each feature, at each timepoint, and in each condition, all features were binned and visualized in the pie chart according to the condition in which that feature achieved its maximum mean intensity across all timepoints. (B) Kendrick mass defect (KMD) correlation of 112 ^12^C/^13^C pairs that were identified by visual inspection of mass spectra. The features were scaled with (14/14.01565) for a ^12^CH_2_ subunit and (15/15.01900) for a ^13^CH_2_ subunit. Linear regression was defined by *y* = 1.056*x* – 0.00199, yielding a correlation coefficient of R^2^ = 0.9995. (C) Van Krevelen analysis showing the hydrogen/carbon (H/C) versus the oxygen/carbon (O/C) ratios for all 112 ^13^C features. The outlined windows represent regions of diagnostic H/C and O/C ratios established by Laszakovits and MacKay to characterize molecules from different classes,^25^ except for the polyphenol window, which was established by Oesterle et al.^26^ (D) Scores plot for principal component analysis of the signal intensity across all timepoints and experiment conditions for the 112 ^13^C features; datapoints depict the distribution of timepoints and colored ellipses represent 95% confidence intervals, with statistics generated via 999 PERMANOVA permutations. (E) Distribution of 112 ^13^C features from the GAM_+fecal/+lignin_ condition, according to size (*m/z*) and the time during incubation at which each achieved its maximum mean intensity; no features reached a maximum intensity at the 72 hour timepoint. See Tables S4–S8 for source data.

An initial assessment of the mass spectra was made with MS-Dial (V5.5.251021) to align chromatographic peaks and define mass spectrometric features according to *m/z* and retention time. A total of 13,234 features were identified (**Table S4**). Each feature was normalized by subtracting the mean signal intensity of the instrument blank and the GAM_-fecal/-lignin_ control samples. Negative or 0 values were imputed with 10 a.u. (a value more than 5 times lower than the lower limit of detection) to avoid computational issues from missing or zero values in quotient or log mathematical transformations performed during statistical analyses; 3,029 features were excluded from additional analysis because all of their entries across the timeseries were 10 a.u., leaving 10,205 features distributed across the GAM_+fecal/+lignin,_ GAM_-fecal/+lignin_, and GAM_+fecal/-lignin_ conditions (**Figure 3A)**. A preliminary comparison of this feature distribution was performed by calculating the mean signal intensity for each feature at each timepoint in each condition. All features were binned according to the condition in which that feature achieved its maximum mean intensity across all timepoints. Human gut microbiota appear to have adapted their metabolic activity in response to supplemented lignin such that the greatest number of metabolite features with the highest mean intensity was found in cultures containing lignin and human gut microbiota (GAM_+fecal/+lignin_): 4,647 of 10,205 features (45.5%) (**Figure 3A**), compared with 3,292 of 10,205 features (32.2%) detected in GAM_+fecal/-lignin_ cultures. The GAM_-fecal/+lignin_ cultures yielded the lowest number of features (2,266 of 10,205, 22.2%). The increased diversity of metabolites that was uniquely afforded upon culturing human gut microbiota with lignin suggests that this dietary polymer can substantially shape the metabolome of human gut bacterial communities.

### Determination of ^12^C/^13^C Pairs and their Compound Class Identities

To assess the extent to which gut bacterial deconstruction of lignin contributes to this metabolic shift, we identified (by manual review of mass spectra) the metabolites detected as pairs of unlabeled (^12^C) and labeled (per-^13^C) isotopologs, yielding 143 ^12^C/^13^C feature pairs (**Table S4**, **Figure S1**). Of these, 112 were identified by MS-Dial for extraction of intensities; intensities for the remaining pairs could not be extracted, likely due to low signal intensity or overlapping chromatographic peaks. We used Kendrick mass defect (KMD) analysis to assess the 112 pairs because it allows for identification of homologous molecules (**Table S5, Figure S2**).^37^ The analysis is performed by rescaling *m/z* values such that molecules that are a pair of ^12^C/^13^C isotopologs will have mathematically equivalent mass defects (i.e., the difference between the nominal integer *m/z* and the scaled observed *m/z*) (**Table S5**). This equivalency enabled high-throughput confirmation of ^12^C/^13^C pairs, as represented by the linear correlation in **Figure 3B**. Identification of ^12^C/^13^C mass spectral pairs allowed for unequivocal identification of the number of carbon atoms in each feature because the observed *m/z* of the ^13^C monoisotopic peak was greater than the observed *m/z* of the ^12^C monoisotopic peak by the number of carbon atoms in the metabolite (**Figure S1**). This *m/z* differential allowed for accurate molecular formula assignments (**Table S4**).^23^ Across the 112 ^12^C/^13^C feature pairs, there were 65 distinct molecular formulas; features with replicated molecular formulas were each observed at distinct retention times, suggesting that they are isomers.

Van Krevelen analysis of the hydrogen/carbon (H/C) versus oxygen/carbon (O/C) ratios in molecules can be used to broadly determine the classes of compounds present in a complex mixture as well as the types of chemical transformations that relate the molecules in the mixture. Laszakovits and MacKay defined a diagnostic window of H/C and O/C ratios for lignin (0.86–1.34 and 0.21–0.44, respectively), finding that the H/C ratios characterize its high aromatic content.^25^ Analysis of the H/C and O/C ratios in the GAM_+fecal/+lignin_ cultures (**Figure 3C**, **Table S6**) revealed that 84 of 112 (75.0%) features with distinct molecular formulas fit within the diagnostic window for lignin. Of the 28 features outside of the lignin diagnostic window, 14 fell into the window that Laszakovits and MacKay defined for the tannin-type polyphenols. Interestingly, 9 of these 14 features include nitrogen. There is precedent for gut microbiota installing nitrogen on polyphenols: ammonia was covalently linked to the polyphenol epigallocatechin gallate by gut microbiota in mice.^38^ Another 2 of the 28 features fit into the window used to characterize lipids, and further inspection of their molecular formulas and their late retention times suggest that they are hydroxylated fatty acids (**Table S4**). Another feature fell within the window used to classify peptides, but the molecular formula for this feature does not contain nitrogen. Laszakovits and MacKay emphasize that there is overlap in the regions that characterize lignin and peptides (**Figure 3C**), which may explain our result for this feature. There were 11 features that did not fall into any one of the categories established by Laszakovits and MacKay, but they are accounted for by the van Krevelen windows established by Oesterle et al. for polyphenols (**Figure 3C**).^26^

### Principal Component Analysis Revealed Distinct Clustering as a Function of Lignin and Human Gut Microbiota

Metabolic deconstruction of lignin was further characterized through comprehensive assessment of all 112 ^13^C *m/z* features. The unique distribution of these features across all timepoints and conditions (GAM_+fecal/+lignin_, GAM_+fecal/-lignin_, and GAM_-fecal/+lignin_) was revealed via principal component analysis.^27^ The resulting scores plot (**Figure 3D, Table S7**) shows differentiation of distinct sample groups as a function of lignin and gut microbiota presence, where each datapoint represents a replicate at a given timepoint. A subsequent PERMANOVA analysis with 999 permutations resulted in an F-value of 347.8, a correlation of determination of 0.8689, and an adjusted *p*-value of 0.001, indicating that the collection of metabolites (in presence and intensity) was unique for each incubation condition. A broad distribution of data points in the plot corresponds to increased changes in the metabolic profile over the course of the incubation, whereas a tight clustering of data points signifies little change in the number and types of molecules over the course of the incubation. Thus, the dispersion observed for the GAM_+fecal/+lignin_ condition indicates a changing metabolic profile over time, while the tight clustering of samples when the lignin is incubated in the absence of gut microbiota (GAM_-fecal/+lignin_) indicates metabolic stability, consistent with its sterility. The proximity of the GAM_+fecal/+lignin_ (purple) and GAM_-fecal/+lignin_ (green) clusters, combined with their distance from the GAM_+fecal/-lignin_ cluster (blue), represents that supplementation of cultures with lignin significantly shifts the chemical milieu, especially upon the introduction of human gut microbiota.

### Per-^13^C Feature Masses Progressively Decrease in Size Over the Course of Incubation

The processive nature of lignin deconstruction was revealed by plotting the *m/z* of each of the 112 ^13^C features in the GAM_+fecal/+lignin_ condition at the timepoint it reached its mean maximum intensity (**Figure 3E**, **Table S8**). Mass windows were defined to account for lignin as well as lignin fragments and deconstruction products across an *m/z* range: 800–1000 Da (tetrameric–hexameric lignin), 600–799.99 Da (trimeric–pentameric lignin), 400–599.99 Da (dimeric–tetrameric lignin), 225–399.99 Da (dimeric lignin), and 50–224.99 Da (monomeric lignin). Before culturing began, 2 features in the 800–1000 Da range were present at their maximum intensity at the 0-hour timepoint. Lignin oligomers in the 600–799.99 Da and 400–599.99 Da windows were abundant in the 0.25 and 2 hour timepoints (suggesting they arose from larger lignin polymers). After only 2 hours of incubation, the distribution shifted to features with smaller *m/z* values as lignin and large lignin oligomers were deconstructed. Such a rapid deconstruction of lignin is on par with the rate achieved by termites^14^ and points to an inherent capacity within the human gut microbiome to break down lignin.

Over the course of the first 8 hours of incubation, the features generally displayed a broad distribution of *m/z* values, encompassing the entire range from 50–1000 Da. The production of smaller metabolites (monomers and dimers) and oligomers (trimers–hexamers) appears concomitant: depolymerization of intact lignin polymer and large lignin oligomers likely proceeds by successive cleavage of individual terminal constituents. By 24 hours of incubation, which corresponds to the median colonic transit time observed by Nandhra et al.,^39^ the distribution of features is marked by a distinctive trend toward smaller metabolites. In particular, no features in the 800–1000 Da window reached their maximum intensity after 4 hours of incubation, and no features in the 600–799.99 Da window reached their maximum intensity after 24 hours, suggesting that the microbial community had reached a steady-state for the deconstruction of lignin and large lignin oligomers between 24 and 48 hours. Moreover, the majority of distinct metabolites were produced within the first 24 hours: only 11 of the 112 features reached their maximum intensity after 24 hours, and all of these were below 600 Da. No features reached their maximum intensity at 72 hours. Taken together, the distribution of per-^13^C mass spectrometric features observed over 96 hours when human gut microbiota were incubated with per-^13^C lignin trends from larger to smaller *m/z* values as intact lignin polymers were deconstructed by human gut microbiota into oligomeric fragments that were subsequently degraded to secondary metabolites.

### Metabolic Profiles Revealed Metabolites are Produced and Consumed

The dynamics of lignin metabolism by human gut microbiota were further detailed by comparing the relative abundance of each of the 112 ^13^C features across all culture conditions and timepoints (**Figure 4A**, **Table S9**). Three qualitative metabolic profiles were readily observed when lignin was incubated with fecal bacteria (GAM_+fecal/+lignin_) as compared to sterile counterparts (GAM_-fecal/+lignin_), termed Consumption, Transient, and Production (**Figures 4B**, **S3**). The Consumption profile is characterized by constant feature intensity in the absence of bacteria; however, upon introduction of fecal bacteria, these features decreased in intensity over time, eventually approaching the minimal intensities observed in the absence of lignin supplementation (GAM_+fecal/-lignin_) (**Figure S3A**). This pattern describes 58 of 112 ^13^C features, with 26 of those being >600 Da, suggesting their presence at the beginning of incubation and subsequent metabolism by human gut microbiota over time (**Tables S9–S10**).

**Figure 4:**
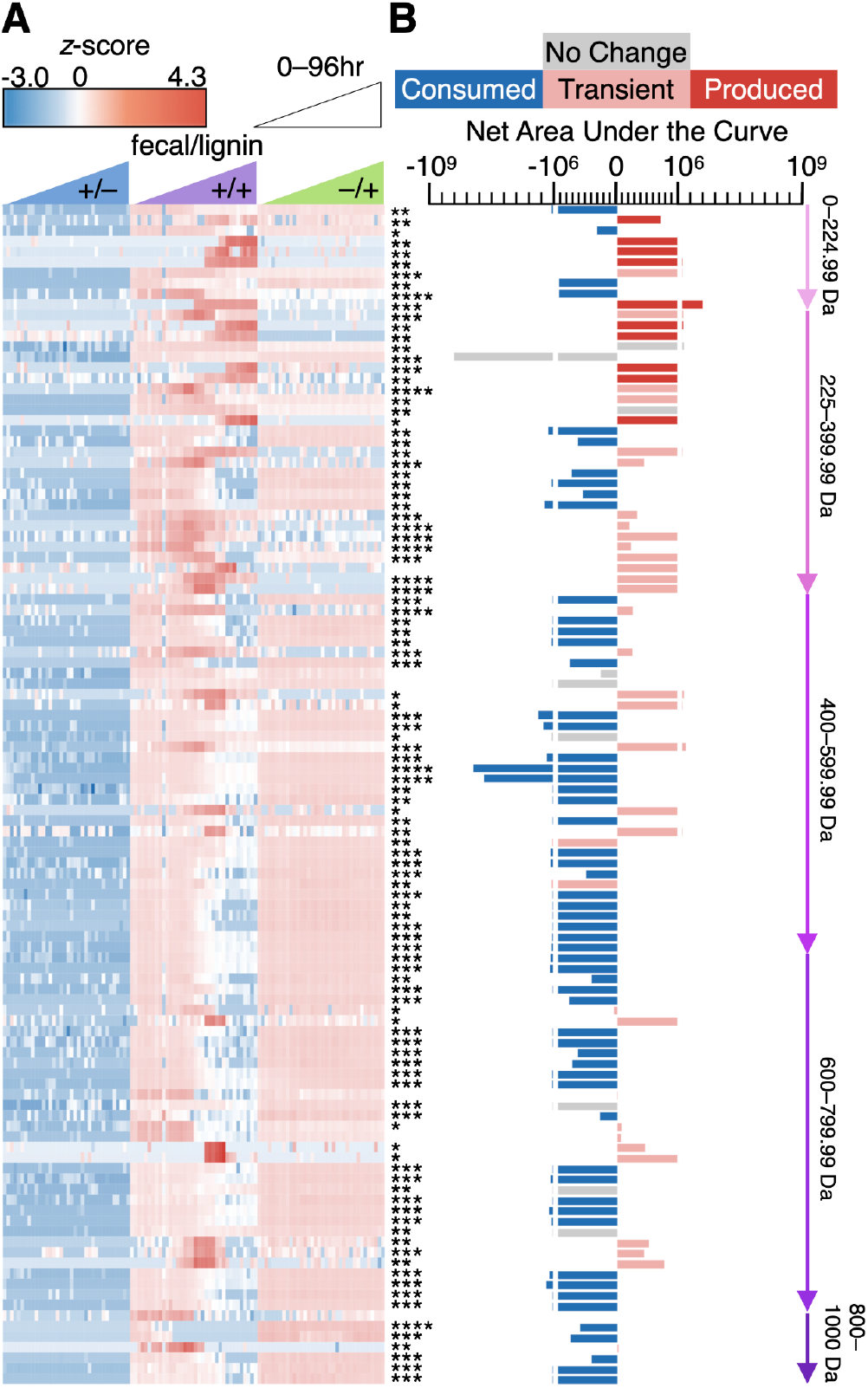
Metabolic profiles and net area under the curve reveal trends that indicate lignin deconstruction. **(A)** Comparison of the signal intensity for 112 per-^13^C features according to sample condition and incubation time, as measured by the *z*-score for signal intensity of each replicate at each timepoint, where red depicts features with increased intensity and blue depicts features with decreased intensity. **(B)** Features are ordered by increasing *m/z*. The bars depict the mean net AUC for each of the 112 ^13^C features, represented as the difference of the mean GAM_+fecal/+lignin_ AUC minus the mean GAM_-fecal/+lignin_ AUC, and the bar colors depict the feature’s metabolic profile. Discovery was assessed by unpaired *t*-test with the two-stage step-up method of Benjamini, Krieger, and Yekutieli multiple testing correction (false discovery rate Q=1%; n = 3; *p*-adjusted: *, <0.05; **, <0.01; ***, <0.001; ****, <0.0001). See **Tables S9** and **S10** for source data.

Consumption profiles were complemented by Transient and Production metabolic profiles because these features showed production only in the presence of both lignin and fecal bacteria. Features with a Transient metabolic profile were produced after the start of the incubation and were subsequently consumed. This profile is typified by the feature with a ^13^C *m/z* of 381.2310 Da, identified as secoisolariciresinol. The position of secoisolariciresinol in the middle of the lignan bioconversion pathway (**Figure 1A**) corroborates its observed Transient metabolic profile (**Figure S3B**).

Features with a Production profile are produced after the start of incubation but, unlike the Transient profile, they persist and show no evidence of subsequent consumption. Such features may represent kinetic or thermodynamic barriers in a biochemical pathway. For example, the feature with a ^13^C *m/z* of 315.1738 Da, identified as enterolactone, displayed a Production metabolic profile (**Figure S3C**), a trait that matches its putative position at the end of the lignan bioconversion pathway (**Figure 1A**).^40^ Together, these metabolic patterns demonstrate that communities of human gut bacteria from fecal samples deconstructed per-^13^C lignin and produced a variety of lignin-derived secondary metabolites.

### AUC of Metabolic Profiles Revealed Progressive Lignin Deconstruction by Human Gut Microbiota

Quantitative comparison of patterns in metabolite profiles was performed by first determining the area under the curve (AUC) for each metabolite across the entire incubation. As shown in **Figure 4B**, of the 112 ^13^C features assessed in the presence of lignin, 103 (92.0%) displayed AUCs significantly altered as a function of gut microbiota presence (i.e., GAM_+fecal/+lignin_ relative to GAM_-fecal/+lignin_). The extent of these differences was calculated as the mean AUC for the GAM_-fecal/+lignin_ control condition subtracted from the mean AUC for the GAM_+fecal/+lignin_ condition (**Equation 4**). Features whose fold change between their maximum mean intensity and minimum mean intensity was less than 10 in the GAM_+fecal/+lignin_ samples were characterized as No Change (9 of 112 features; **Figure S3D**, **Table S10**). Features displaying a Consumption profile yielded a net negative AUC, while metabolites with Production profiles yielded a net positive AUC (**Figure 4B**). The net AUC values for features characterized by a Transient profile were predominantly positive (31 of 35), while 4 Transient features had net negative AUC values (**Table S10**).

The existence of both Consumption and Transient profiles in the 800–1000 Da feature class indicates that lignin and large lignin oligomers are deconstructed by human gut microbiota (**Table S10**). Of the 7 features in this window, 6 had a net negative AUC value and displayed a Consumption profile. These large lignin oligomers were present at the beginning of the incubation, and they were deconstructed over time in the presence of human gut microbiota. More striking is the remaining feature in this mass range because it displays a Transient profile with a net positive AUC value. This lignin oligomer was not present at the beginning of incubation. Instead, it arose over the course of incubation, putatively via deconstruction of lignin polymer larger than 1000 Da, only to be subsequently deconstructed itself.

The features in the 600–799.99 Da range are also characterized by the Consumption and Transient metabolic profiles. The 21 features with net negative AUC values and Consumption profiles reinforce the finding that human gut microbiota deconstruct lignin oligomers. The progressive deconstruction of lignin oligomers to smaller fragments is shown by the 10 features in this mass range with Transient profiles: as larger oligomers are broken down, the features in this range are temporarily produced, only to be metabolized themselves.

The features in the 400–599.99 Da range serve to link the deconstruction of larger features to the production of smaller features. While 22 of 34 features in this range display Consumption profiles, 9 features display Transient profiles and 3 are characterized as No Change. The range from 225–399.99 Da marked the shift toward the Production profile for the smaller features: as lignin and lignin oligomers were deconstructed, their dimeric and monomeric constituents were produced. Only 6 of the 27 features in this window were characterized by a Consumption profile, while 13 were characterized by a Transient profile and 5 were characterized by Production profiles; the remaining 3 features in this range were characterized as No Change. Similarly, only 4 of 10 features in the 50–224.99 Da range displayed a Consumption profile, while 1 was characterized as Transient and 5 were characterized by Production profiles, demonstrating that the monomeric metabolites generally resulted from the deconstruction of lignin.

The metabolic profiles of features that result when human gut microbiota are incubated with lignin display a trend that leads from the degradation of larger features toward the production of smaller features over time. This trend characterizes the deconstruction of lignin and the metabolism of the resulting oligomers by human gut microbiota, leading to the production of dimeric and monomeric polyphenols. Overall, these results demonstrate, for the first time, that human gut microbiota deconstruct intact lignin polymer and large lignin oligomers, and they demonstrate that the process is rapid, occurring on the scale of hours.

### Mass Spectrometry Reveals Putative Structures of a Network of Lignin Deconstruction Metabolites

We next sought to identify plausible structures for the features with the largest *m/z* as well as for their metabolic products (**Figure 5**). With evidence for a large lignin oligomer containing 47 carbon atoms (**Table S4**), we examined mass spectrometry data to assemble the structure of lignin oligomer **A**. The mass spectrum of **A** (**Figure S4A.3**) showed the mirrored isotope pattern that characterizes a ^12^C/^13^C pair, indicating its origin in lignin supplemented to cultures. This metabolite is characterized by a Consumption profile (**Figure S4A.4**), consistent with a large oligomer that is metabolized over time. Although the signal intensity for this feature was insufficient to obtain fragmentation spectra, likely due to poor solubility and ion suppression,^41^ we observed that a 17-carbon fragment was common to the fragmentation spectra of many detected ^12^C/^13^C pairs of metabolites (**Figure S4**). We hypothesized that this fragment was a core structural subunit of lignin oligomer **A**. We identified this subunit as tricin (C_17_H_14_O_7_) because it contains 17 carbon atoms, (**Figure 5A**), it was detected in samples via NMR (**Figure 2**), and it is a common end-group of wheat straw lignin, the source of lignin in these experiments.^34^

**Figure 5:**
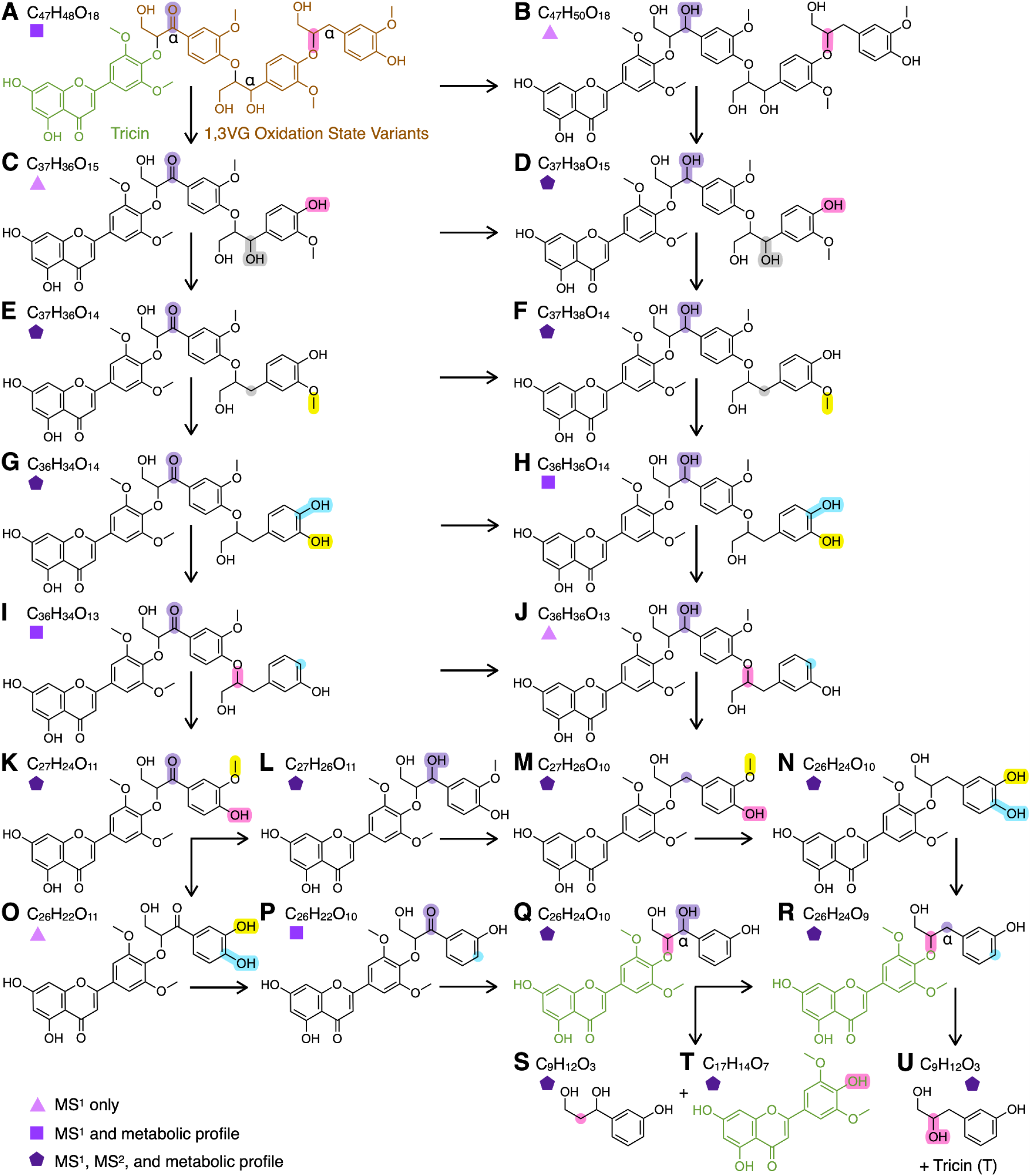
A large network of metabolites is produced when human gut microbiota deconstruct lignin. (A–U) Structures of 21 metabolites related to the deconstruction of lignin. Human gut microbiota perform a series of reductions, hydrolyses, demethylations, and dehydroxylations to transform lignin oligomer **A** into tricin (metabolite **T**, colored green) and phenolic monomers, metabolites **S** and **U**. Highlighted bonds and atoms track the transformations from one metabolite to the next. Pink highlights depict the β-O-4 bonds and phenolic cleavage products. Brown coloring in **A** depicts (3-(4-hydroxy-3-methoxyphenyl)propan-1,3-diol (1,3VG). Cɑ positions are labelled in metabolites **A**, **Q,** and **R**. Colored shapes represent the strength of the structural identification, where structures with light purple triangles are characterized by MS^1^ mass spectra only; structures with medium purple squares are characterized by MS^1^ mass spectra and a full metabolic profile; and structures with dark purple pentagons are characterized by MS^1^ mass spectra, MS^2^ fragmentation mass spectra, and full metabolic profiles (**Figure S4**).

Accounting for the other 30 carbons in lignin oligomer **A**, we hypothesized three oxidation state variants of another common lignin subunit: a positional isomer of vanillyl glycol, 3-(4-hydroxy-3-methoxyphenyl)propan-1,3-diol (10 carbon atoms, putatively C_10_H_13_O_4_, termed 1,3VG) (**Figure 5A, Figure S4D.6**).^42,43,44^ The following variants are supported by the mass spectrum of **A** and by putative metabolites of its deconstruction. In one variant, the Cɑ position was oxidized as a carbonyl. In another, the carbonyl was reduced to a hydroxyl group. In the third, the Cɑ position was fully reduced. We hypothesized that these three 1,3VG variants and tricin were linked to afford lignin oligomer **A**.

To test our hypothesized structure of lignin oligomer **A**, we queried our panel of identified ^12^C/^13^C pairs for metabolites that would derive from **A**’s deconstruction by human gut microbiota. The result was 20 metabolites interconnected in a biochemical network by dehydroxylation, demethylation, hydrolysis, and reduction—all reaction classes performed by human gut microbiota (**Figure 5B–U**).^45,46^ Metabolite **B** was identified as a product of carbonyl reduction of **A**. Depolymerization of metabolite **B** produced metabolite **D**, whose overlaid ^12^C and ^13^C fragmentation spectra indicate the presence of fragments corresponding to metabolites **L**, **T**, and 1,3VG **(Figure S4D.5**–**S4D.6)**. The finding that larger lignin oligomers were comprised of smaller metabolites supports our putative structure of **A** and its deconstruction. Moreover, 13 of the metabolites in this network produced viable fragmentation spectra, 9 of which showed spectra consistent with tricin (metabolite **T**) as a common fragment (**Figure S4**).

Proceeding along a different metabolic route, after depolymerization of lignin oligomer **A**, dehydroxylation of the ɑ-hydroxy group on metabolite **C** could afford metabolite **E**. Demethylation of metabolite **E** produced metabolite **G**, whose Transient metabolic profile **(Figure S4G.4)** is consistent with a metabolite that is first produced upon lignin’s incubation with gut bacteria before being degraded itself into smaller metabolites. The overlaid ^12^C and ^13^C fragmentation spectra for metabolite **G** (**Figure S4G.5**) yielded *m/z* values of 493.1202 Da and 519.2043 Da, corresponding to the respective ^12^C and ^13^C *m/z* values for metabolite **P**, substantiating **G** within this metabolic network. Subsequent dehydroxylation of metabolite **G** produced metabolite **I**. The biochemical transformation of guaiacyl subunits by demethylation of the 3-methoxy moiety followed by dehydroxylation of the 4-OH group are well-characterized activities of human gut microbiota,^32^ occurring in many transformations throughout this metabolic network.

Depolymerization, demethylation, and dehydroxylation led from metabolite **I** to metabolite **P**. Reduction of the carbonyl to a secondary alcohol formed metabolite **Q**, followed by dehydroxylation to produce metabolite **R**. The overlaid ^12^C and ^13^C fragmentation spectra of metabolite **R** (**Figure S4R.5)** yielded *m/z* values of 329.0660 Da and 346.1227 Da, corresponding to the respective ^12^C and ^13^C *m/z* values for tricin, metabolite **T**. The metabolic profile for tricin (**Figure S4T.4**) was observed as No Change because the fold-change between its maximum and minimum intensities in the GAM_+fecal/+lignin_ condition was less than 10, a stringent threshold applied to avoid over-representing ambiguous cases as being metabolized by human gut microbiota (**Table S4**). Unbound tricin appears to be present in very high quantities in the lignin substrate (green shading, GAM_-fecal/+lignin_), and it appears to be metabolized quickly by human gut microbiota (purple shading), dropping to its lowest level by 8 hours. However, tricin levels appear to rebound after 12 hours and maintain a relatively steady state for the remainder of the 96 hour incubation. In the same way that enterolactone production is explained by sustained deconstruction of lignin feeding into the enterolignan biosynthetic pathway (**Figure 1**), sustained depolymerization of lignin and lignin oligomers to the tricin end-group explains its sustained presence throughout the duration of the incubation.

In addition to yielding tricin, depolymerization of metabolite **R** by cleavage of the β-O-4 linkage produced metabolite **U**, characterized by a Production profile (**Figure S4U.4**). The fragmentation spectra for metabolite **U** show neutral loss of formaldehyde from the per-^13^C isotopolog, consistent with fragmentation patterns of 1,2-diols (**Figure S4U.5**).^47^

### Evidence for Lignin Depolymerization by Human Gut Microbiota

Metabolomic analyses suggested that lignin is depolymerized by cleavage of the β-O-4 linkages and that this depolymerization occurs in two different ways, depending on the substrate (**Figure S5**). Metabolite **U** lacked a hydroxy group at the Cɑ position, which appears to result in hydrolysis of the β-O-4 bond. However, when the hydroxy group was present in the Cɑ position (**Figure 5Q**), then depolymerization of the β-O-4 linkage appears to occur via reduction. Inspection of the fragmentation spectra for metabolite **S**, a positional isomer of metabolite **U**, showed neutral loss of ethanol (**Figure S5E**), consistent with a 1,3-diol. Neutral loss of formaldehyde was not seen in the fragmentation spectra of metabolite **S**, suggesting that the hydroxyl group is in a different position from that of metabolite **U**.

The mechanism by which cleavage of the β-O-4 bond occurs under anaerobic conditions remains unknown, but evidence from enzymes in soil bacteria provides a rational starting point. An entire family of aerobic β-ether reductase enzymes found in the soil bacterium *Sphingomonas paucimobilis* has been characterized for the reduction of the β-O-4 bond in lignin.^48^ Some of these enzymes cleave the β-O-4 bonds between subunits with a hydroxy group in the Cɑ position, which is consistent with our proposed lignin oligomer structures.^49^ Moreover, facultative anaerobic soil bacteria that contain β-ether reductase genes homologous to those in *S. paucimobilis* have been identified,^50^ supporting the feasibility of anaerobic lignin depolymerization by different types of soil bacteria.^6^ This was an important finding because anaerobic deconstruction of lignin has historically been considered unlikely. Our search for homologous lignin-degrading β-ether reductases in human gut microbiotas yielded no matches with significant homology, yet the existence of such enzymes in anaerobic soil organisms support the biochemical plausibility of metabolism via this route in the human gut.

Collectively, our data show that human gut microbiota deconstruct lignin under anaerobic conditions, and we identified putative structures of large lignin oligomers that are deconstructed into smaller metabolites across an extensive metabolic network. While only single-stage mass spectrometric spectra are available for the largest lignin oligomers and a few downstream metabolites in the network, likely due to issues with solubility and ionization, these findings provide a foundation for subsequent characterization via mass spectrometric fragmentation spectra and NMR analyses to strengthen the veracity of our proposed structures. We have assembled evidence that suggests that deconstruction of the plausible metabolites in this network involves anaerobic cleavage of the β-O-4 ether linkage in lignin, providing motivation for future characterization of the specific human gut bacteria and enzymes responsible for this process.

## Conclusion

For over a century, the ability of human gut microbiota to digest lignin has been debated.^2,3,12,13^ At the core of this tension was a challenge of analytical chemistry: definitively characterizing lignin and tracing the diverse array of metabolites that can arise from metabolism of this irregular, heterogeneous polymer. Our initial handle for tracking metabolic flux downstream of lignin was provided by application of chemical logic. Namely, because pinoresinol is a sub-structure embedded in the lignin polymer, we hypothesized that it may be abstracted from lignin and subsequently fed into a known multi-step, multi-bacteria biochemical pathway^32,33^ resulting in enterolactone as the final product. Using per-^13^C lignin—which enabled us to distinguish lignin and its metabolites from the metabolome endogenous to fecal samples from individuals consuming dietary lignin—we revealed that sustained deconstruction of lignin by human gut microbiota releases resinol-type lignans (e.g., pinoresinol), feeding continual production of enterolactone. Additionally, we proposed the structure of a large lignin oligomer, finding that its metabolic deconstruction by human gut microbiota spawned a network of 20 additional metabolites, some of which resulted from depolymerization of the β-O-4 ether bonds. Complementary 2D NMR analyses supported these findings: a reduction in the relative abundance of β-O-4 ether bonds—the most common linkage in lignin of wheat straw—represented microbial cleavage of the lignin polymer.

Enterolactone deriving from lignin is translationally significant. Numerous studies in cell culture,^51^ rodents,^30^ and humans show that enterolactone can decrease incidence^52^ and risk^53^ of hormone-dependent breast cancer.^54^ Our findings suggest that consuming lignin fiber may be a vehicle to elevate enterolactone levels. Given that cancer prevention requires long-term intervention,^55^ diet modification is a strategy with one of the lowest economic barriers to success.^56^

Beyond targeted analysis, untargeted LC–MS analyses provided a holistic view of lignin deconstruction dynamics. From the array of 112 ^12^C/^13^C metabolite pairs revealed and tracked over time, a shift from large oligomers that predominated at the start of culture (Consumption profile) to smaller metabolites that became more prevalent as the culture continued (Production profile) demonstrated a processive metabolism of lignin. Further compelling is the Transient profile of large lignin oligomers, which appeared following the start of culturing and subsequently disappeared. These mid-culture pulses of lignin oligomers indicated continuous deconstruction of lignin polymers that are larger than the mass spectrometry upper limit of detection configured for our experiments. Targeted analysis of mass spectrometric fragments enabled the structural identification of one such large lignin oligomer and the monitoring of its deconstruction over time.

Among these metabolites, we found flavonoids like tricin, whose anti-inflammatory effects can help decrease the risk of chronic conditions such as cardiomyopathy^57^ and type-2 diabetes.^58^ We found enterolignans, whose phytoestrogenic character has been associated with a decreased risk of breast cancer. Future work will aim to identify the bacteria responsible for lignin deconstruction and to characterize the breadth of phenolic metabolites that are produced, including from lignin sourced from other plants that each have a unique polymer composition.^4^ Our work contributes to the development of a new paradigm by establishing that the human gut microbiota contains an inherent capacity to deconstruct lignin—even in anaerobic conditions—a phenomenon historically considered impossible.

## AUTHOR INFORMATION

### Author Contributions

CAD, DJY, and ENB designed experiments. CAD, DJY, and ENB conducted data analysis. CAD, EAS, DJY, and JDR performed experiments. CAD, DJY, and ENB wrote and edited the manuscript. ENB acquired funds. ENB provided project supervision and administration. All authors read and approved the final version of the manuscript.

### Note

The authors declare no competing financial interests.

### Data Availability

Data for this article, including quantitative LC–MS/MS data, untargeted LC–MS data, and targeted LC–MS/MS fragmentation data, are available at Metabolomics Workbench (Project IDs 8332 and 8333).

## ACKNOWLEDGEMENTS

This study was made possible in part through access to the Mass Spectrometry Core Facility of the Department of Chemistry, a shared resource at the University of California, Irvine. We appreciate the contributions of facility director Dr. Felix Grün and proteomics specialist Ben Katz, who provided technical assistance and experimental guidance. We also appreciate the Fraunhofer Center for Chemical-Biotechnological Processes generously providing us with organosolv wheat lignin. The table of contents graphic was created with BioRender.com (agreement number JM2AAMMSO9).

## TOC Graphic

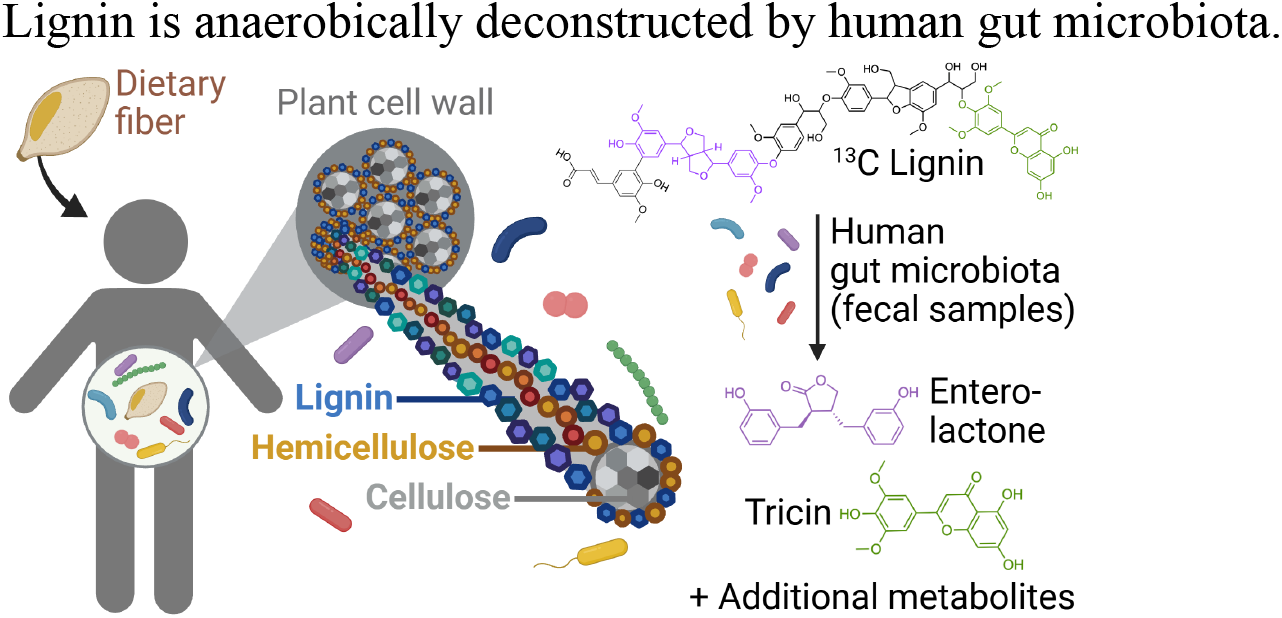

## REFERENCES

1. National, F. I. National Food Institute, Technical University of Denmark. Frida Food Data. Frida Fooddata.Dk. Https://Frida.Fooddata.Dk.

(2) Williams, R. D.; Olmsted, W. H.; Hamann, C. H.; Fiorito, J. A.; Duckles, D. The Effect of Cellulose, Hemicellulose and Lignin on the Weight of the Stool: A Contribution to the Study of Laxation in Man. The Journal of Nutrition 1936, 11 (5), 433–449. 10.1093/jn/11.5.433

(3) Kelsay, J. L.; Goering, H. K.; Behall, K. M.; Prather, E. S. Effect of Fiber from Fruits and Vegetables on Metabolic Responses of Human Subjects: Fiber Intakes, Fecal Excretions, and Apparent Digestibilities. The American Journal of Clinical Nutrition 1981, 34 (9), 1849–1852. 10.1093/ajcn/34.9.1849

(4) Boerjan, W.; Ralph, J.; Baucher, M. Lignin Biosynthesis. Annu. Rev. Plant Biol. 2003, 54 (1), 519–546. 10.1146/annurev.arplant.54.031902.134938

(5) Vanholme, R.; Demedts, B.; Morreel, K.; Ralph, J.; Boerjan, W. Lignin Biosynthesis and Structure. Plant Physiology 2010, 153 (3), 895–905. 10.1104/pp.110.155119

(6) DeAngelis, K. M.; Sharma, D.; Varney, R.; Simmons, B.; Isern, N. G.; Markilllie, L. M.; Nicora, C.; Norbeck, A. D.; Taylor, R. C.; Aldrich, J. T.; Robinson, E. W. Evidence Supporting Dissimilatory and Assimilatory Lignin Degradation in *Enterobacter Lignolyticus* SCF1. Front. Microbiol. 2013, 4. 10.3389/fmicb.2013.00280

(7) Chaput, G.; Ford, J.; DeDiego, L.; Narayanan, A.; Tam, W. Y.; Whalen, M.; Huntemann, M.; Clum, A.; Spunde, A.; Pillay, M.; Palaniappan, K.; Varghese, N.; Mikhailova, N.; Chen, I.M.; Stamatis, D.; Reddy, T. B. K.; O’Malley, R.; Daum, C.; Shapiro, N.; Ivanova, N.; Kyrpides, N. C.; Woyke, T.; Glavina Del Rio, T.; DeAngelis, K. M. *Sodalis Ligni* Strain 159R Isolated from an Anaerobic Lignin-Degrading Consortium. Microbiol. Spectr. 2022, 10 (3), e02346-21. 10.1128/spectrum.02346-21

(8) Brown, M. E.; Chang, M. C. Exploring Bacterial Lignin Degradation. Current Opinion in Chemical Biology 2014, 19, 1–7. 10.1016/j.cbpa.2013.11.015

(9) Tuor, U.; Winterhalter, K.; Fiechter, A. Enzymes of White-Rot Fungi Involved in Lignin Degradation and Ecological Determinants for Wood Decay. Journal of Biotechnology 1995, 41 (1), 1–17. 10.1016/0168-1656(95)00042-O

(10) Tien, M.; Kirk, T. K. Lignin-Degrading Enzyme from the Hymenomycete *Phanerochaete Chrysosporium* Burds. Science 1983, 221 (4611), 661–663. 10.1126/science.221.4611.661

(11) Janusz, G.; Pawlik, A.; Sulej, J.; Świderska-Burek, U.; Jarosz-Wilkołazka, A.; Paszczyński, A. Lignin Degradation: Microorganisms, Enzymes Involved, Genomes Analysis and Evolution. FEMS Microbiology Reviews 2017, 41 (6), 941–962. 10.1093/femsre/fux049

(12) Kirk, T. K.; Farrell, R. L. ENZYMATIC “COMBUSTION”: THE MICROBIAL DEGRADATION OF LIGNIN. Ann. Rev. Microbiol. 1987, No. 41, 465–505.

13. Asp, N.-G. Dietary Fibre-Definition, Chemistry and Analytical Determination. Molecular Aspects of Medicine 1987, 9 (1), 17–29. 10.1016/0098-2997(87)90014-8

14. Li, H.; Yelle, D. J.; Li, C.; Yang, M.; Ke, J.; Zhang, R.; Liu, Y.; Zhu, N.; Liang, S.; Mo, X.; Ralph, J.; Currie, C. R.; Mo, J. Lignocellulose Pretreatment in a Fungus-Cultivating Termite. Proc. Natl. Acad. Sci. U.S.A. 2017, 114 (18), 4709–4714. 10.1073/pnas.1618360114

(15) Begum, A. N.; Nicolle, C.; Rémésy, C.; Scalbert, A.; Mila, I.; Lapierre, C.; Nagano, K.; Fukushima, K.; Heinonen, S.M.; Adlercreutz, H. Dietary Lignins Are Precursors of Mammalian Lignans in Rats. The Journal of Nutrition 2004, 134 (1), 120–127. 10.1093/jn/134.1.120

(16) Lankiewicz, T. S.; Choudhary, H.; Gao, Y.; Amer, B.; Lillington, S. P.; Leggieri, P. A.; Brown, J. L.; Swift, C. L.; Lipzen, A.; Na, H.; Amirebrahimi, M.; Theodorou, M. K.; Baidoo, E. E. K.; Barry, K.; Grigoriev, I. V.; Timokhin, V. I.; Gladden, J.; Singh, S.; Mortimer, J. C.; Ralph, J.; Simmons, B. A.; Singer, S. W.; O’Malley, M. A. Lignin Deconstruction by Anaerobic Fungi. Nat. Microbiol. 2023, 8 (4), 596–610. 10.1038/s41564-023-01336-8

(17) Kasimir, M.; Behrens, M.; Humpf, H.U. Release of Small Phenolic Metabolites from Isotopically Labeled ^13^C Lignin in the Pig Cecum Model. J. Agric. Food Chem. 2022, 70 (27), 8317–8325. 10.1021/acs.jafc.2c02836

(18) Niemi, P.; Aura, A.M.; Maukonen, J.; Smeds, A. I.; Mattila, I.; Niemelä, K.; Tamminen, T.; Faulds, C. B.; Buchert, J.; Poutanen, K. Interactions of a Lignin-Rich Fraction from Brewer’s Spent Grain with Gut Microbiota in Vitro. J. Agric. Food Chem. 2013, 61 (27), 6754–6762. 10.1021/jf401738x

(19) Aura, A.M.; Niemi, P.; Mattila, I.; Niemelä, K.; Smeds, A.; Tamminen, T.; Faulds, C.; Buchert, J.; Poutanen, K. Release of Small Phenolic Compounds from Brewer’s Spent Grain and Its Lignin Fractions by Human Intestinal Microbiota in Vitro. J. Agric. Food Chem. 2013, 61 (40), 9744–9753. 10.1021/jf4024195

(20) Dicksion, C. A.; Chao, D. N.; Rickmeyer, J. D.; Bess, E. N. A Validated LC–MS/MS Method for Quantifying Phenolic Acids, Lignans, and Enterolignans from Human Fecal Samples. ACS Nutr. Sci. 2026, 1 (4), 356–366. 10.1021/acsnutrsci.6c00063

(21) Yelle, D. J.; Ralph, J.; Frihart, C. R. Characterization of Nonderivatized Plant Cell Walls Using High-resolution Solution-state NMR Spectroscopy. Magnetic Reson. in Chemistry 2008, 46 (6), 508–517. 10.1002/mrc.2201

(22) Wang, F.; Allen, D.; Tian, S.; Oler, E.; Gautam, V.; Greiner, R.; Metz, T. O.; Wishart, D. S. CFM-ID 4.0 – a Web Server for Accurate MS-Based Metabolite Identification. Nucleic Acids Research 2022, 50 (W1), W165–W174. 10.1093/nar/gkac383

(23) Patiny, L.; Borel, A. ChemCalc: A Building Block for Tomorrow’s Chemical Infrastructure. J. Chem. Inf. Model. 2013, 53 (5), 1223–1228. 10.1021/ci300563h

(24) Krutkin, D. D.; Thomas, S.; Zuffa, S.; Rajkumar, P.; Knight, R.; Dorrestein, P. C.; Kelley, S. T. To Impute or Not To Impute in Untargeted Metabolomics─That Is the Compositional Question. J. Am. Soc. Mass Spectrom. 2025, 36 (4), 742–759. 10.1021/jasms.4c00434

(25) Laszakovits, J. R.; MacKay, A. A. Data-Based Chemical Class Regions for Van Krevelen Diagrams. J. Am. Soc. Mass Spectrom. 2022, 33 (1), 198–202. 10.1021/jasms.1c00230

(26) Oesterle, I.; Pristner, M.; Berger, S.; Wang, M.; Verri Hernandes, V.; Rompel, A.; Warth, B. Exposomic Biomonitoring of Polyphenols by Non-Targeted Analysis and Suspect Screening. Anal. Chem. 2023, 95 (28), 10686–10694. 10.1021/acs.analchem.3c01393

(27) Pang, Z.; Lu, Y.; Zhou, G.; Hui, F.; Xu, L.; Viau, C.; Spigelman, A. F.; MacDonald, P. E.; Wishart, D. S.; Li, S.; Xia, J. MetaboAnalyst 6.0: Towards a Unified Platform for Metabolomics Data Processing, Analysis and Interpretation. Nucleic Acids Research 2024, 52 (W1), W398–W406. 10.1093/nar/gkae253

(28) Zeng, J.; Helms, G. L.; Gao, X.; Chen, S. Quantification of Wheat Straw Lignin Structure by Comprehensive NMR Analysis. J. Agric. Food Chem. 2013, 61 (46), 10848–10857. 10.1021/jf4030486

(29) Heinonen, S.; Nurmi, T.; Liukkonen, K.; Poutanen, K.; Wähälä, K.; Deyama, T.; Nishibe, S.; Adlercreutz, H. In Vitro Metabolism of Plant Lignans: New Precursors of Mammalian Lignans Enterolactone and Enterodiol. J. Agric. Food. Chem. 2001, 49 (7), 3178–3186. 10.1021/jf010038a

(30) Mabrok, H. B.; Klopfleisch, R.; Ghanem, K. Z.; Clavel, T.; Blaut, M.; Loh, G. Lignan Transformation by Gut Bacteria Lowers Tumor Burden in a Gnotobiotic Rat Model of Breast Cancer. Carcinogenesis 2012, 33 (1), 203–208. 10.1093/carcin/bgr256

(31) Gorissen, A.; Kraut, N. U.; De Visser, R.; De Vries, M.; Roelofsen, H.; Vonk, R. J. No de Novo Sulforaphane Biosynthesis in Broccoli Seedlings. Food Chemistry 2011, 127 (1), 192–196. 10.1016/j.foodchem.2010.12.067

(32) Bess, E. N.; Bisanz, J. E.; Yarza, F.; Bustion, A.; Rich, B. E.; Li, X.; Kitamura, S.; Waligurski, E.; Ang, Q. Y.; Alba, D. L.; Spanogiannopoulos, P.; Nayfach, S.; Koliwad, S. K.; Wolan, D. W.; Franke, A. A.; Turnbaugh, P. J. Genetic Basis for the Cooperative Bioactivation of Plant Lignans by *Eggerthella Lenta* and Other Human Gut Bacteria. Nat. Microbiol. 2020, 5 (1), 56–66. 10.1038/s41564-019-0596-1

(33) Clavel, T.; Henderson, G.; Alpert, C.A.; Philippe, C.; Rigottier-Gois, L.; Doré, J.; Blaut, M. Intestinal Bacterial Communities That Produce Active Estrogen-like Compounds Enterodiol and Enterolactone in Humans. Appl. Environ. Microbiol. 2005, 71 (10), 6077–6085. 10.1128/AEM.71.10.6077-6085.2005

(34) Del Río, J. C.; Rencoret, J.; Prinsen, P.; Martínez, Á. T.; Ralph, J.; Gutiérrez, A. Structural Characterization of Wheat Straw Lignin as Revealed by Analytical Pyrolysis, 2D-NMR, and Reductive Cleavage Methods. J. Agric. Food Chem. 2012, 60 (23), 5922–5935. 10.1021/jf301002n

(35) Rich, B. E.; Jackson, J. C.; De Ora, L. O.; Long, Z. G.; Uyeda, K. S.; Bess, E. N. Alternative Pathway for Dopamine Production by Acetogenic Gut Bacteria That O-Demethylate 3-Methoxytyramine, a Metabolite of Catechol O-Methyltransferase. Journal of Applied Microbiology 2022, 133 (3), 1697–1708. 10.1111/jam.15682

(36) Gall, D. L.; Kontur, W. S.; Lan, W.; Kim, H.; Li, Y.; Ralph, J.; Donohue, T. J.; Noguera, D. R. *In Vitro* Enzymatic Depolymerization of Lignin with Release of Syringyl, Guaiacyl, and Tricin Units. Appl. Environ. Microbiol. 2018, 84 (3), e02076–17. 10.1128/AEM.02076-17

(37) Kendrick, E. A Mass Scale Based on CH_2_ = 14.0000 for High Resolution Mass Spectrometry of Organic Compounds. Anal. Chem. 1963, 35 (13), 2146–2154. 10.1021/ac60206a048

(38) Zhang, S.; Zhao, Y.; Ohland, C.; Jobin, C.; Sang, S. Microbiota Facilitates the Formation of the Aminated Metabolite of Green Tea Polyphenol (-)-Epigallocatechin-3-Gallate Which Trap Deleterious Reactive Endogenous Metabolites. Free Radical Biology and Medicine 2019, 131, 332–344. 10.1016/j.freeradbiomed.2018.12.023

(39) Nandhra, G. K.; Mark, E. B.; Di Tanna, G. L.; Haase, A.; Poulsen, J.; Christodoulides, S.; Kung, V.; Klinge, M. W.; Knudsen, K.; Borghammer, P.; Andersen, K. O.; Fynne, L.; Sutter, N.; Schlageter, V.; Krogh, K.; Drewes, A. M.; Birch, M.; Scott, S. M. Normative Values for Region-specific Colonic and Gastrointestinal Transit Times in 111 Healthy Volunteers Using the 3D-Transit Electromagnet Tracking System: Influence of Age, Gender, and Body Mass Index. Neurogastroenterology Motil. 2020, 32 (2), e13734. 10.1111/nmo.13734

(40) Borriello, S. P.; Setchell, K. D. R.; Axelson, M.; Lawson, A. M. Production and Metabolism of Lignans by the Human Faecal Flora. Journal of Applied Bacteriology 1985, 58 (1), 37–43. 10.1111/j.1365-2672.1985.tb01427.x

(41) Alseekh, S.; Aharoni, A.; Brotman, Y.; Contrepois, K.; D’Auria, J.; Ewald, J.; C. Ewald, J.; Fraser, P. D.; Giavalisco, P.; Hall, R. D.; Heinemann, M.; Link, H.; Luo, J.; Neumann, S.; Nielsen, J.; Perez De Souza, L.; Saito, K.; Sauer, U.; Schroeder, F. C.; Schuster, S.; Siuzdak, G.; Skirycz, A.; Sumner, L. W.; Snyder, M. P.; Tang, H.; Tohge, T.; Wang, Y.; Wen, W.; Wu, S.; Xu, G.; Zamboni, N.; Fernie, A. R. Mass Spectrometry-Based Metabolomics: A Guide for Annotation, Quantification and Best Reporting Practices. Nat. Methods 2021, 18 (7), 747–756. 10.1038/s41592-021-01197-1

(42) Morreel, K.; Dima, O.; Kim, H.; Lu, F.; Niculaes, C.; Vanholme, R.; Dauwe, R.; Goeminne, G.; Inzé, D.; Messens, E.; Ralph, J.; Boerjan, W. Mass Spectrometry-Based Sequencing of Lignin Oligomers. Plant Physiol. 2010, 153 (4), 1464–1478. 10.1104/pp.110.156489

(43) Huis, R.; Morreel, K.; Fliniaux, O.; Lucau-Danila, A.; Fénart, S.; Grec, S.; Neutelings, G.; Chabbert, B.; Mesnard, F.; Boerjan, W.; Hawkins, S. Natural Hypolignification is Associated with Extensive Oligolignol Accumulation in Flax Stems. Plant Physiology 2012, 158 (4), 1893–1915. 10.1104/pp.111.192328

(44) Kiyota, E.; Mazzafera, P.; Sawaya, A. C. H. F. Analysis of Soluble Lignin in Sugarcane by Ultrahigh Performance Liquid Chromatography–Tandem Mass Spectrometry with a Do-It-Yourself Oligomer Database. Anal. Chem. 2012, 84 (16), 7015–7020. 10.1021/ac301112y

(45) Spanogiannopoulos, P.; Bess, E. N.; Carmody, R. N.; Turnbaugh, P. J. The Microbial Pharmacists within Us: A Metagenomic View of Xenobiotic Metabolism. Nat. Rev. Microbiol. 2016, 14 (5), 273–287. 10.1038/nrmicro.2016.17

(46) Koppel, N.; Maini Rekdal, V.; Balskus, E. P. Chemical Transformation of Xenobiotics by the Human Gut Microbiota. Science 2017, 356 (6344), eaag2770. 10.1126/science.aag2770

(47) McLafferty, F.W; Turecek, F. Interpretation of Mass Spectra, 4th Ed. (McLafferty, Fred W.; Turecek, Frantisek). J. Chem. Educ. 1994, 71 (2), A54. 10.1021/ed071pA54.5

(48) Masai, E.; Ichimura, A.; Sato, Y.; Miyauchi, K.; Katayama, Y.; Fukuda, M. Roles of the Enantioselective Glutathione *S* -Transferases in Cleavage of β-Aryl Ether. J. Bacteriol. 2003, 185 (6), 1768–1775. 10.1128/JB.185.6.1768-1775.2003

(49) Sonoki, T.; Iimura, Y.; Masai, E.; Kajita, S.; Katayama, Y. Specific Degradation of β-Aryl Ether Linkage in Synthetic Lignin (Dehydrogenative Polymerizate) by Bacterial Enzymes of *Sphingomonas paucimobilis* SYK-6 Produced in Recombinant *Escherichia coli*. J. Wood Sci. 2002, 48, 429–433. 10.1007/BF00770705

(50) Billings, A. F.; Fortney, J. L.; Hazen, T. C.; Simmons, B.; Davenport, K. W.; Goodwin, L.; Ivanova, N.; Kyrpides, N. C.; Mavromatis, K.; Woyke, T.; DeAngelis, K. M. Genome Sequence and Description of the Anaerobic Lignin-Degrading Bacterium *Tolumonas lignolytica* Sp. Nov. Stand. in Genomic Sci. 2015, 10 (1), 106. 10.1186/s40793-015-0100-3

(51) Brooks, J. D.; Thompson, L. U. Mammalian Lignans and Genistein Decrease the Activities of Aromatase and 17beta-Hydroxysteroid Dehydrogenase in MCF-7 Cells. J. Steroid Biochem. Mol. Biol, 2005, 94 (5), 461–467. 10.1016/j.jsbmb.2005.02.002

(52) Hu, Y.; Li, Y.; Sampson, L.; Wang, M.; Manson, J. E.; Rimm, E.; Sun, Q. Lignan Intake and Risk of Coronary Heart Disease. Journal of the American College of Cardiology 2021, 78 (7), 666–678. 10.1016/j.jacc.2021.05.049

(53) Adlercreutz, H.; Fotsis, T.; Heikkinen, R.; Dwyer, J. T.; Woods, M.; Goldin, B. R.; Gorbach, S. L. Excretion of the Lignans Enterolactone and Enterodiol and of Equol in Omnivorous and Vegetarian Postmenopausal Women and in Women with Breast Cancer. Lancet 1982, 2 (8311), 1295–1299. 10.1016/s0140-6736(82)91507-0

(54) Adlercreutz, H. Phyto-Oestrogens and Cancer. The Lancet Oncology 2002, 3 (6), 364–373. 10.1016/S1470-2045(02)00777-5

(55) Early Breast Cancer Trialists’ Collaborative Group (EBCTCG). Effects of Chemotherapy and Hormonal Therapy for Early Breast Cancer on Recurrence and 15-Year Survival: An Overview of the Randomised Trials. The Lancet 2005, 365 (9472), 1687–1717. 10.1016/S0140-6736(05)66544-0

(56) Mittelman, S. D. The Role of Diet in Cancer Prevention and Chemotherapy Efficacy. Annu. Rev. Nutr. 2020, 40 (1), 273–297. 10.1146/annurev-nutr-013120-041149

(57) Yu, R.; Zhang, Y.; Wang, T.; Duan, J.; Li, X. Effect of Tricin on Cardiomyocyte Damage Caused by Diabetic Cardiomyopathy (DCM). BMC Cardiovasc. Disord. 2024, 24 (1), 668. 10.1186/s12872-024-04295-y

(58) Chavanelle, V.; F. Otero, Y.; Fayat Cazaubiel, M.; Bard, J.-M.; Pereira, B.; Sapone, V.; Bouchard-Mercier, A.; Bargetto, M.; Boisseau, N.; Guigas, B.; Peltier, S.; Le Joubioux, F.; Maugard, T.; Hadjadj, S.; Sirvent, P.; Marette, A. Totum-63, a Plant-Based Polyphenol-Rich Extract, in Prediabetes or Early-Stage Type 2 Diabetes: An International, Multicenter, Randomized Controlled Trial. Nat. Commun. 2026, 17 (1), 8844. 10.1038/s41467-026-75626-0

