## Supplemental Information for "Metabolomic Investigations Reveal Lignin Deconstruction by Human Gut Microbiota"

**Table S1: Mass Spectrometric Parameters for Quantitation via LC–MS/MS.** Mass spectrometric parameters for the selective quantitation of analytes in enterolignan biosynthetic pathway. Natural isotope abundance and uniformly $^{13}C$-labelled (per-$^{13}C$) isotopologs, along with internal standards, were analyzed with multi-reaction monitoring in negative-ion mode. Q1 and Q3 are the precursor and fragment ions' *m/z*, respectively. CV is the cone voltage, and CE is the collision energy.

| | Q1 (m/z) | Q3 (m/z) | CV (V) | CE (eV) |
|---|---|---|---|---|
| **Lignans** | | | | |
| Lariciresinol (LARI) | 359.0 | 329.0 | 10 | 10 |
| per-$^{13}C$ LARI | 379.0 | 348.0 | 10 | 10 |
| Pinoresinol (PINO) | 357.0 | 151.0 | 20 | 20 |
| per-$^{13}C$ PINO | 377.0 | 159.0 | 20 | 20 |
| Secoisolariciresinol (SECO) | 361.0 | 165.0 | 20 | 20 |
| per-$^{13}C$ SECO | 381.0 | 174.0 | 20 | 20 |
| $d_6$-Secoisolariciresinol | 367.2 | 167.9 | 20 | 30 |
| **Enterolignans** | | | | |
| Enterodiol (END) | 301.0 | 253.0 | 10 | 25 |
| per-$^{13}C$ END | 319.0 | 270.0 | 10 | 25 |
| Enterolactone (ENL) | 297.0 | 107.0 | 20 | 30 |
| per-$^{13}C$ ENL | 315.0 | 114.0 | 20 | 30 |
| $^{13}C_3$-Enterolactone | 300.0 | 254.9 | 20 | 20 |

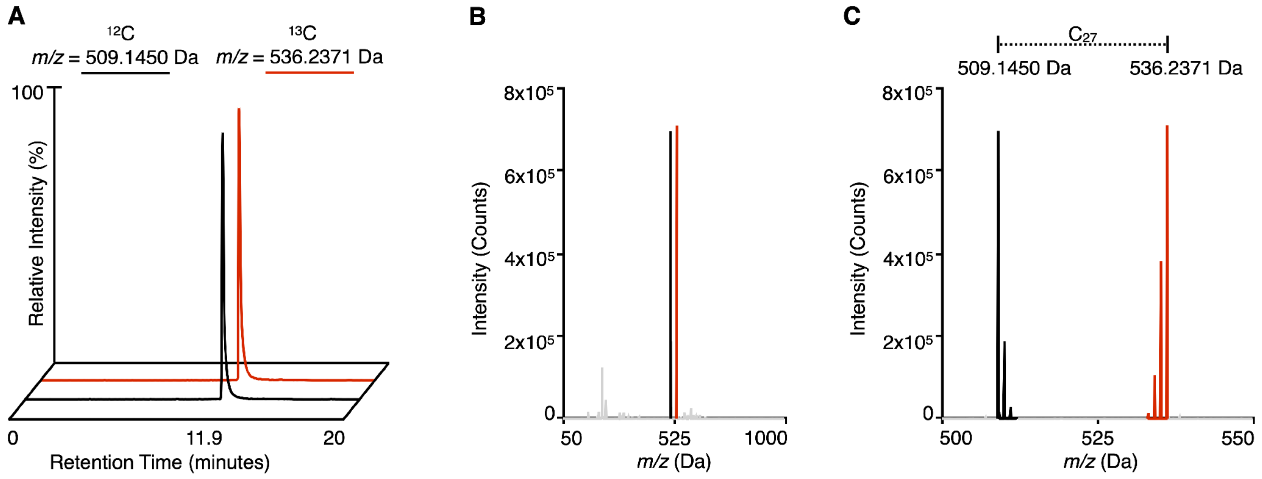


**Figure S1: Representative chromatograms and mass spectra for manually curated $^{12}$C/$^{13}$C pairs of lignin-derived metabolites. (A)** Overlaid extracted ion chromatograms for a $^{12}$C (black) and $^{13}$C (red) feature pair to emphasize co-elution. **(B)** MS$^1$ mass spectrum, showing the full *m/z* range collected (50–1000 Da), extracted from a total ion chromatogram at 11.87 minutes. **(C)** Zoomed in display of the mass spectrum presented in **(B)** to emphasize the mirror-like isotope patterns of $^{12}$C/$^{13}$C pairs. The difference in the nominal masses of the monoisotopic peaks of the pair is used to determine the number of carbon atoms in the metabolite that is represented by the pair.

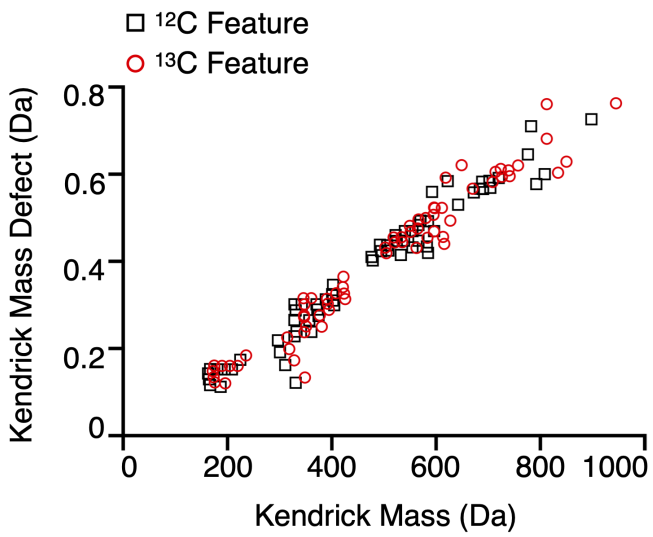


**Figure S2: Overlay of Kendrick mass defect versus Kendrick mass for 112 $^{12}C$ and $^{13}C$ features identified by visual inspection of mass spectra.** A plot of the Kendrick mass defect versus the Kendrick mass can be used to spot isotopologs in a complex mixture. The features were scaled using 14/14.01565 for a $^{12}CH_2$ subunit and 15/15.01900 for a $^{13}CH_2$ subunit to identify the Kendrick Mass. The overlay of the data points for the $^{12}C$ (black) and $^{13}C$ (red) features shows that the features identified as $^{12}C$ and $^{13}C$ pairs display identical distributions on the plot.

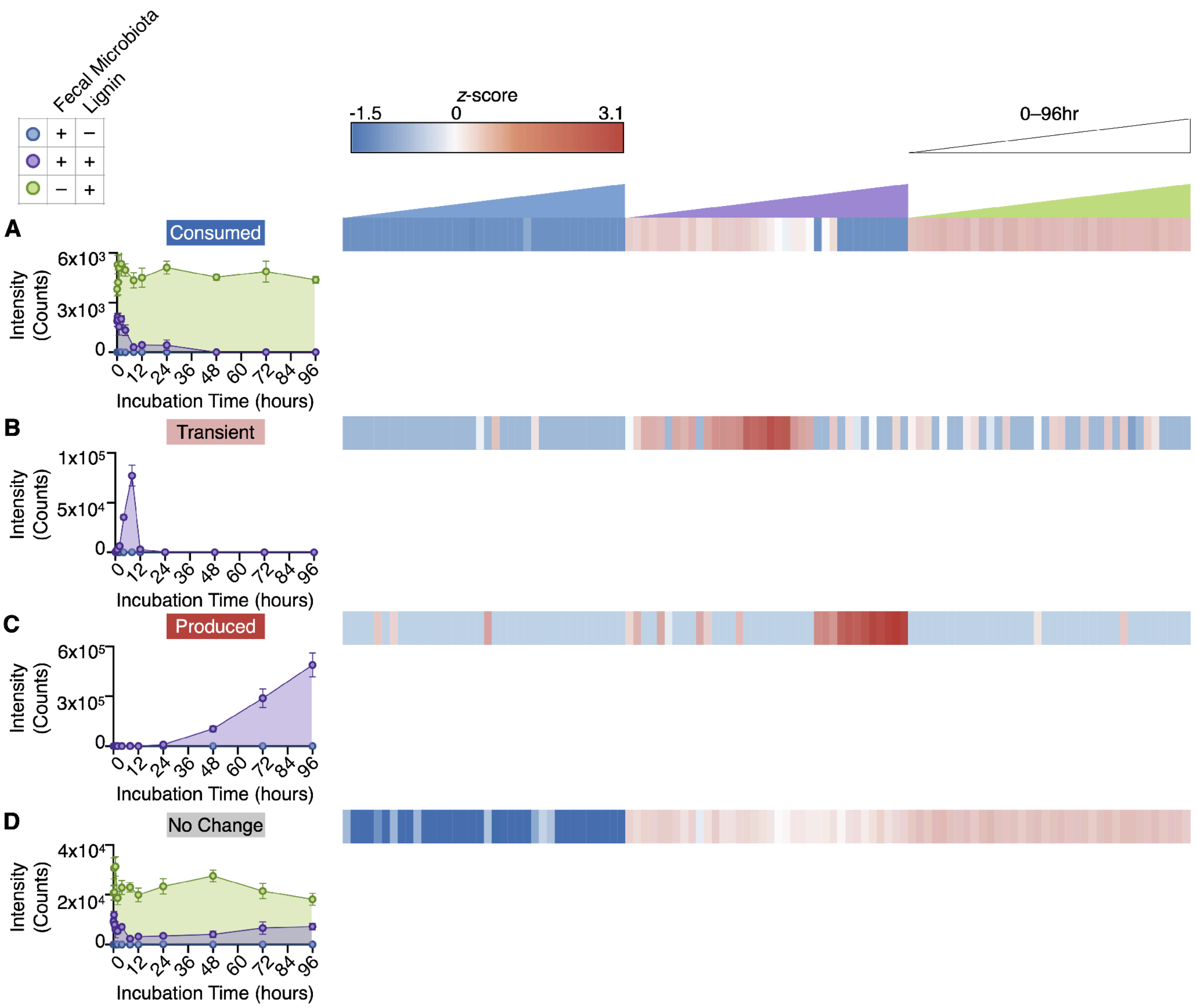


**Figure S3: Representative metabolic profiles of per-$^{13}$C features. (A–D)** Overlays of signal intensity versus incubation time depict metabolic profiles and area under the curve (AUC) for the $GAM_{+fecal/-lignin}$ (blue), $GAM_{+fecal/+lignin}$ (purple), and $GAM_{-fecal/+lignin}$ (green) conditions for **(A)** Consumption metabolic profile (*m/z* = 851.4482 Da, elution time = 10.78 minutes), **(B)** Transient metabolic profile (*m/z* = 381.2322 Da corresponding to secoisolariciresinol, elution time = 6.35 minutes) and **(C)** Production metabolic profile (*m/z* = 315.1732 Da corresponding to enterolactone, elution time = 8.94 minutes). **(D)** No Change metabolic profile (*m/z* = 740.3275 Da, elution time = 8.54 minutes). Data points and AUC depict mean values, and error bars depict the standard error of the mean (n = 3). Heatmap depicts representative metabolic profiles, quantified by the *z*-score for the signal intensity of each replicate (n = 3) at each incubation time-point across all conditions.

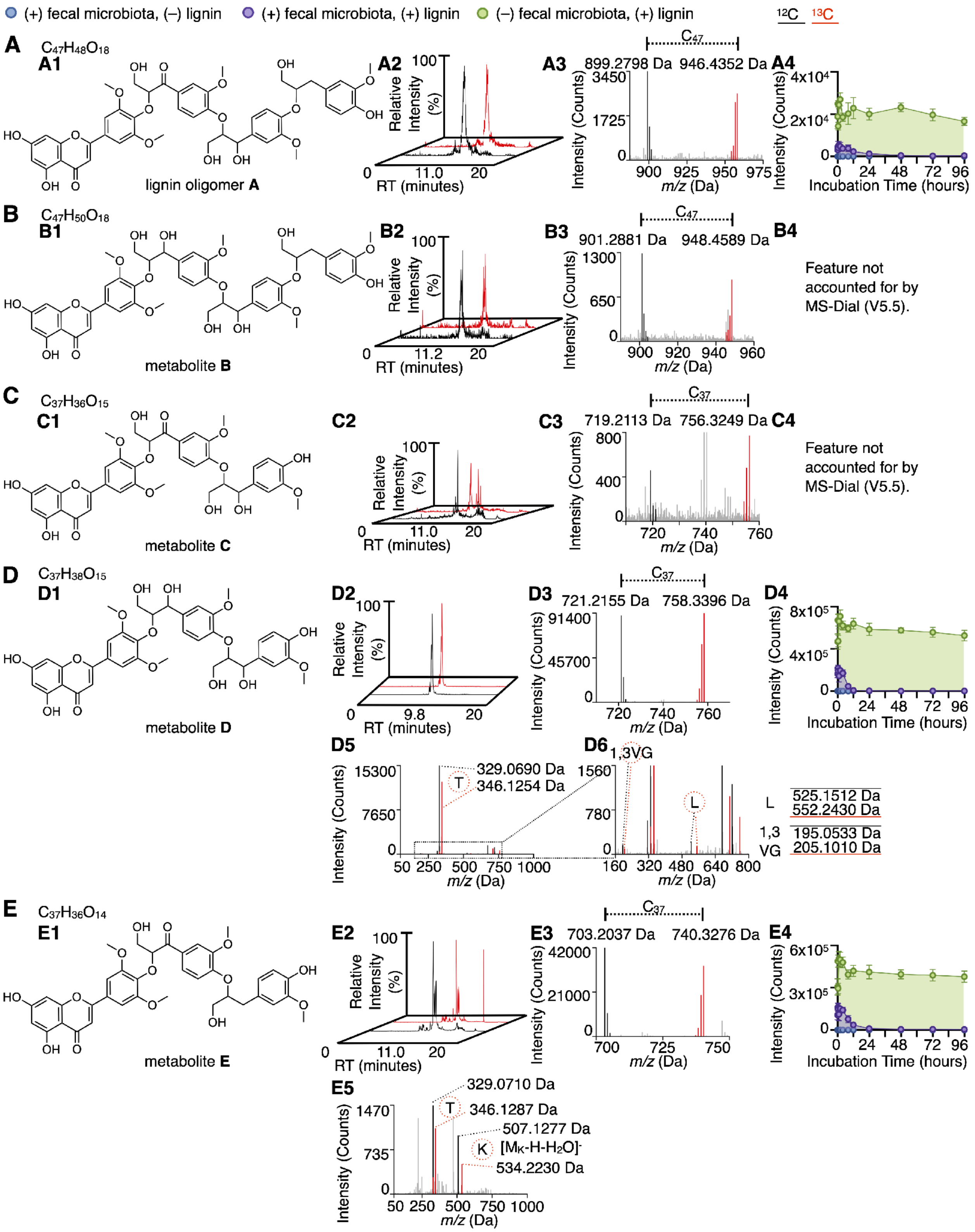

(+) fecal microbiota, (–) lignin
(+) fecal microbiota, (+) lignin
(–) fecal microbiota, (+) lignin
12C 13C
A
C47H48O18
A1
lignin oligomer A
A2
Relative Intensity (%)
RT (minutes)
11.0
A3
C47
899.2798 Da
946.4352 Da
Intensity (Counts)
m/z (Da)
A4
Intensity (Counts)
Incubation Time (hours)
B
C47H50O18
B1
metabolite B
B2
11.2
B3
C47
901.2881 Da
948.4589 Da
B4
Feature not accounted for by MS-Dial (V5.5).
C
C37H36O15
C1
metabolite C
C2
C3
C37
719.2113 Da
756.3249 Da
C4
Feature not accounted for by MS-Dial (V5.5).
D
C37H38O15
D1
metabolite D
D2
9.8
D3
C37
721.2155 Da
758.3396 Da
D4
D5
329.0690 Da
T
346.1254 Da
D6
1,3VG
L
525.1512 Da
552.2430 Da
1,3 VG
195.0533 Da
205.1010 Da
E
C37H36O14
E1
metabolite E
E2
E3
C37
703.2037 Da
740.3276 Da
E4
E5
329.0710 Da
346.1287 Da
507.1277 Da
K
[MK-H-H2O]-
534.2230 Da

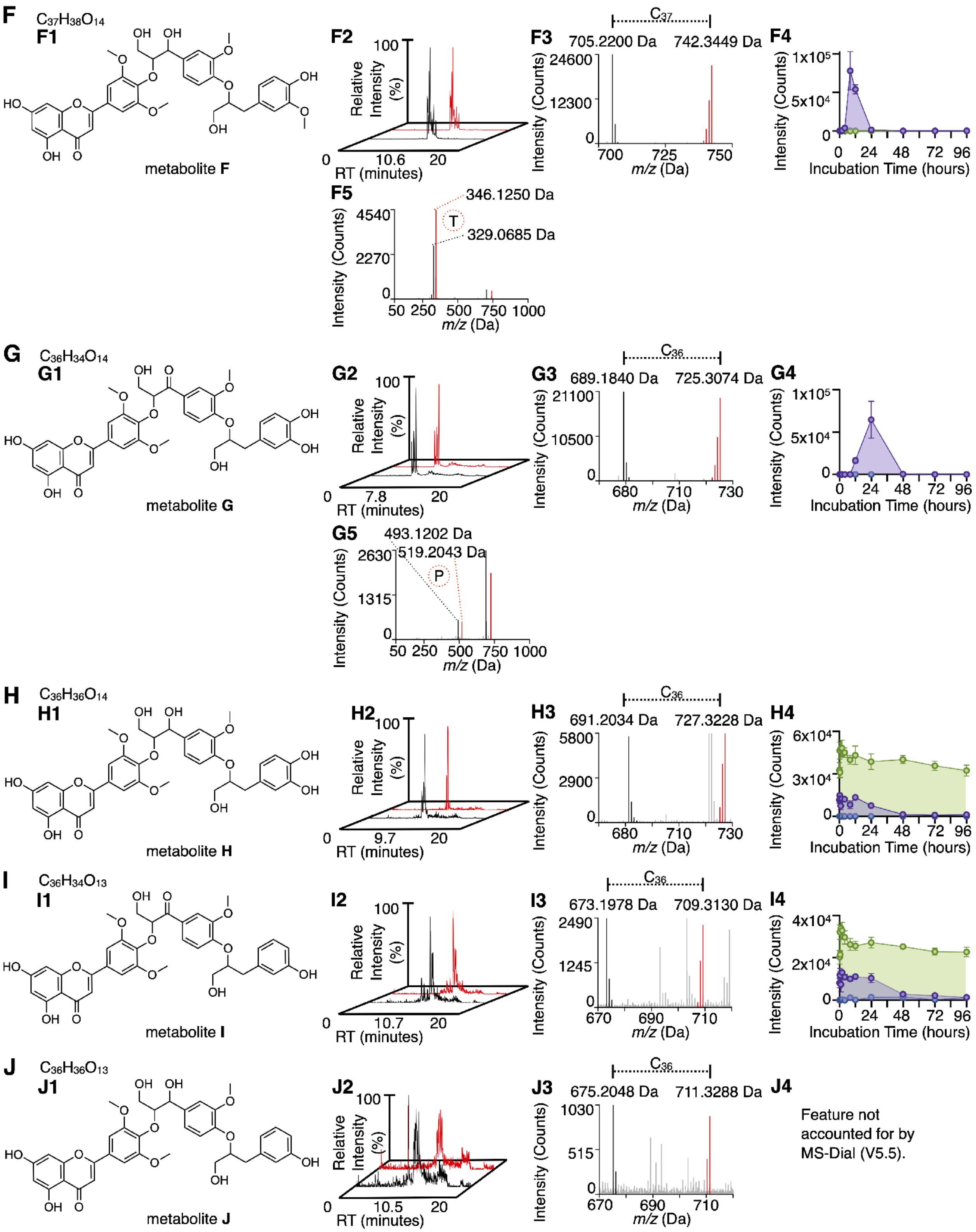
F
$C_{37}H_{38}O_{14}$
F1
metabolite F
F2
Relative Intensity (%)
100
0
10.6
20
RT (minutes)
F3
C37
705.2200 Da
742.3449 Da
Intensity (Counts)
24600
12300
0
700
725
750
m/z (Da)
F4
Intensity (Counts)
1x10^5
5x10^4
0
0 24 48 72 96
Incubation Time (hours)
F5
346.1250 Da
T
329.0685 Da
Intensity (Counts)
4540
2270
0
50 250 500 750 1000
m/z (Da)
G
$C_{36}H_{34}O_{14}$
G1
metabolite G
G2
Relative Intensity (%)
100
0
7.8
20
RT (minutes)
G3
C36
689.1840 Da
725.3074 Da
Intensity (Counts)
21100
10500
0
680
710
730
m/z (Da)
G4
Intensity (Counts)
1x10^5
5x10^4
0
0 24 48 72 96
Incubation Time (hours)
G5
493.1202 Da
519.2043 Da
P
Intensity (Counts)
2630
1315
0
50 250 500 750 1000
m/z (Da)
H
$C_{36}H_{36}O_{14}$
H1
metabolite H
H2
Relative Intensity (%)
100
0
9.7
20
RT (minutes)
H3
C36
691.2034 Da
727.3228 Da
Intensity (Counts)
5800
2900
0
680
710
730
m/z (Da)
H4
Intensity (Counts)
6x10^4
3x10^4
0
0 24 48 72 96
Incubation Time (hours)
I
$C_{36}H_{34}O_{13}$
I1
metabolite I
I2
Relative Intensity (%)
100
0
10.7
20
RT (minutes)
I3
C36
673.1978 Da
709.3130 Da
Intensity (Counts)
2490
1245
0
670
690
710
m/z (Da)
I4
Intensity (Counts)
4x10^4
2x10^4
0
0 24 48 72 96
Incubation Time (hours)
J
$C_{36}H_{36}O_{13}$
J1
metabolite J
J2
Relative Intensity (%)
100
0
10.5
20
RT (minutes)
J3
C36
675.2048 Da
711.3288 Da
Intensity (Counts)
1030
515
0
670
690
710
m/z (Da)
J4
Feature not accounted for by MS-Dial (V5.5).

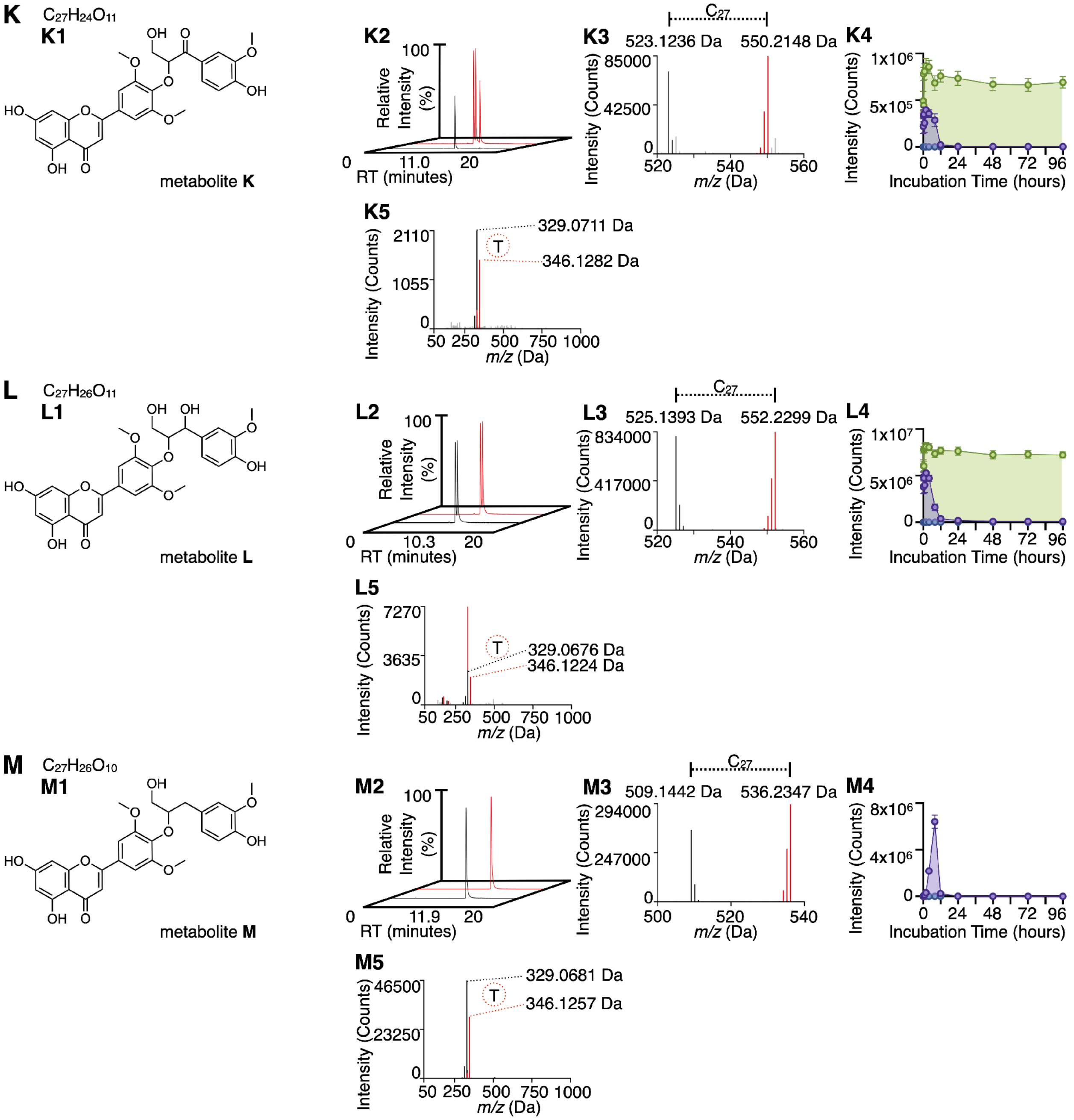
K
C27H24O11
K1
metabolite K
K2
Relative Intensity (%)
100
0
11.0
20
RT (minutes)
K3
C27
523.1236 Da
550.2148 Da
Intensity (Counts)
85000
42500
0
520
540
560
m/z (Da)
K4
1x10^6
5x10^5
0
24
48
72
96
Incubation Time (hours)
K5
329.0711 Da
T
346.1282 Da
2110
1055
0
50 250 500 750 1000
m/z (Da)
L
C27H26O11
L1
metabolite L
L2
100
0
10.3
20
RT (minutes)
L3
C27
525.1393 Da
552.2299 Da
834000
417000
0
520
540
560
m/z (Da)
L4
1x10^7
5x10^6
0
24
48
72
96
Incubation Time (hours)
L5
7270
3635
0
T
329.0676 Da
346.1224 Da
50 250 500 750 1000
m/z (Da)
M
C27H26O10
M1
metabolite M
M2
100
0
11.9
20
RT (minutes)
M3
C27
509.1442 Da
536.2347 Da
294000
247000
0
500
520
540
m/z (Da)
M4
8x10^6
4x10^6
0
24
48
72
96
Incubation Time (hours)
M5
329.0681 Da
T
346.1257 Da
46500
23250
0
50 250 500 750 1000
m/z (Da)

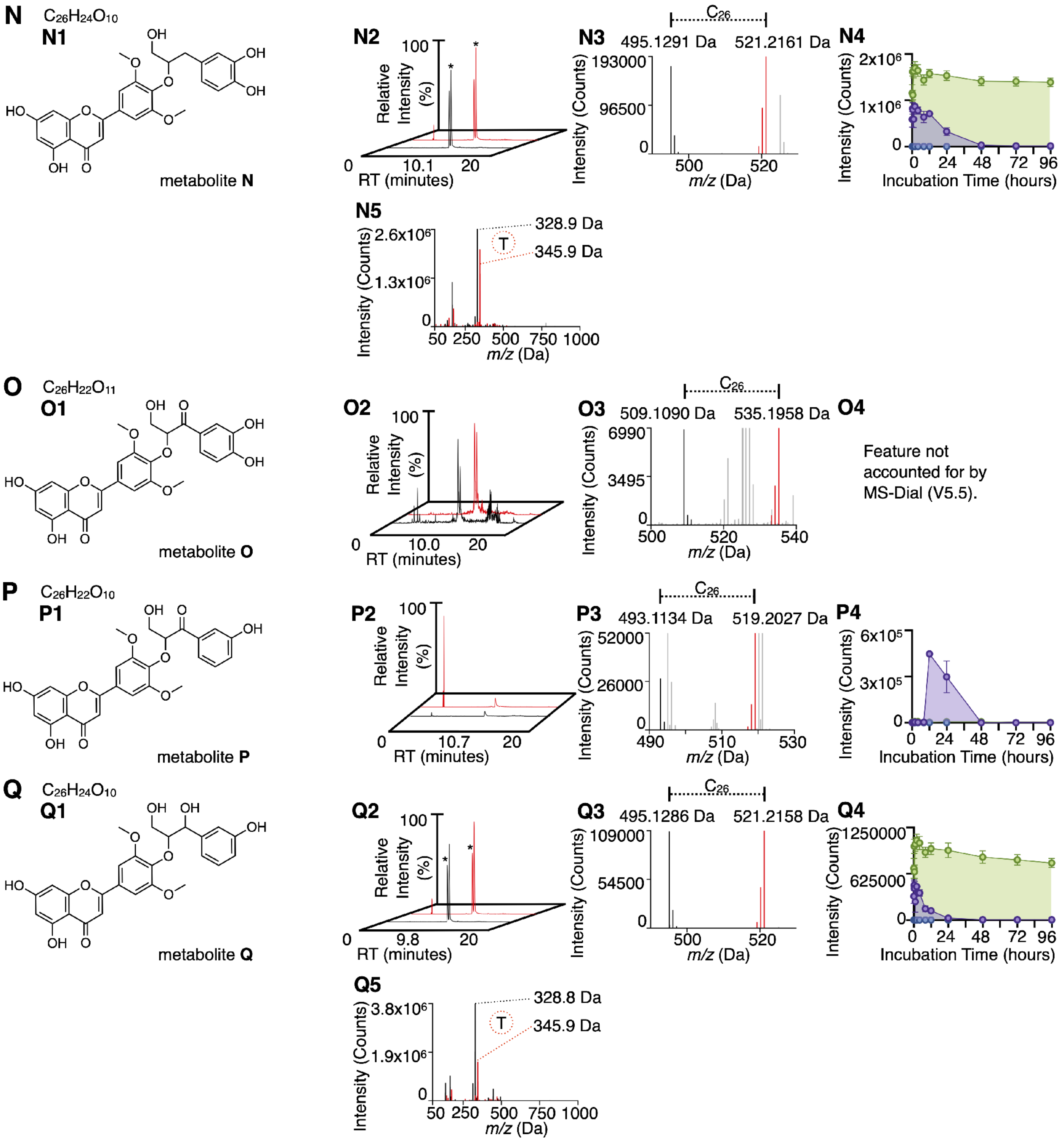
N
C26H24O10
N1
metabolite N
N2
Relative Intensity (%)
100
0
10.1
20
RT (minutes)
N3
C26
495.1291 Da
521.2161 Da
Intensity (Counts)
193000
96500
0
500
520
m/z (Da)
N4
Intensity (Counts)
2x10⁶
1x10⁶
0
0 24 48 72 96
Incubation Time (hours)
N5
Intensity (Counts)
2.6x10⁶
1.3x10⁶
0
328.9 Da
345.9 Da
T
50 250 500 750 1000
m/z (Da)
O
C26H22O11
O1
metabolite O
O2
Relative Intensity (%)
100
0
10.0
20
RT (minutes)
O3
C26
509.1090 Da
535.1958 Da
Intensity (Counts)
6990
3495
0
500
520
540
m/z (Da)
O4
Feature not accounted for by MS-Dial (V5.5).
P
C26H22O10
P1
metabolite P
P2
Relative Intensity (%)
100
0
10.7
20
RT (minutes)
P3
C26
493.1134 Da
519.2027 Da
Intensity (Counts)
52000
26000
0
490
510
530
m/z (Da)
P4
Intensity (Counts)
6x10⁵
3x10⁵
0
0 24 48 72 96
Incubation Time (hours)
Q
C26H24O10
Q1
metabolite Q
Q2
Relative Intensity (%)
100
0
9.8
20
RT (minutes)
Q3
C26
495.1286 Da
521.2158 Da
Intensity (Counts)
109000
54500
0
500
520
m/z (Da)
Q4
Intensity (Counts)
1250000
625000
0
0 24 48 72 96
Incubation Time (hours)
Q5
Intensity (Counts)
3.8x10⁶
1.9x10⁶
0
328.8 Da
345.9 Da
T
50 250 500 750 1000
m/z (Da)

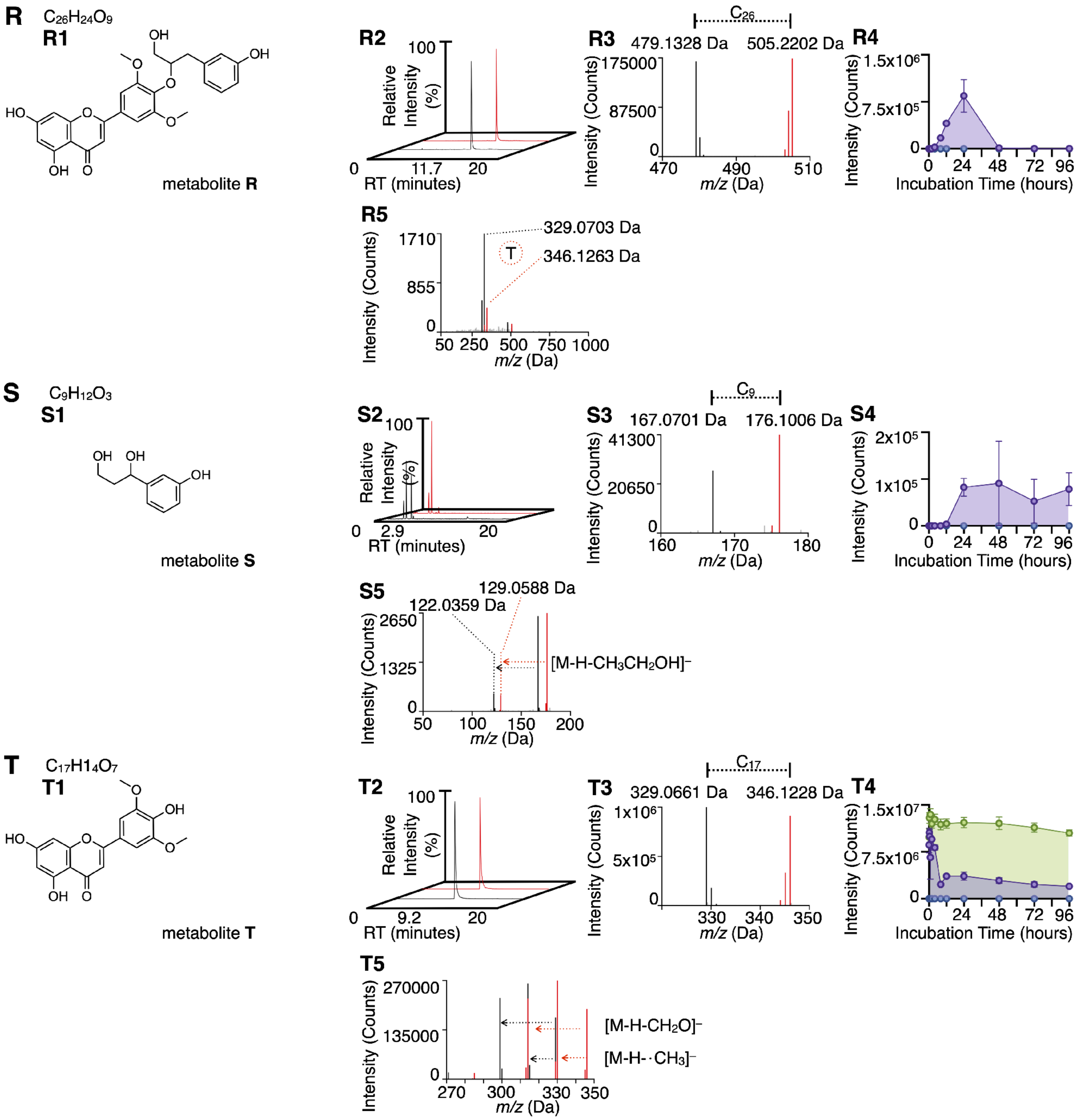

R
C26H24O9
R1
OH
OH
O
O
HO
O
O
OH
O
metabolite R
R2
100
Relative Intensity (%)
0
11.7
20
RT (minutes)
R3
C26
479.1328 Da
505.2202 Da
175000
87500
0
Intensity (Counts)
470
490
510
m/z (Da)
R4
1.5x10^6
7.5x10^5
0
Intensity (Counts)
0
24
48
72
96
Incubation Time (hours)
R5
1710
855
0
Intensity (Counts)
329.0703 Da
346.1263 Da
T
50
250
500
750
1000
m/z (Da)
S
C9H12O3
S1
OH
OH
OH
metabolite S
S2
100
Relative Intensity (%)
0
2.9
20
RT (minutes)
S3
C9
167.0701 Da
176.1006 Da
41300
20650
0
Intensity (Counts)
160
170
180
m/z (Da)
S4
2x10^5
1x10^5
0
Intensity (Counts)
0
24
48
72
96
Incubation Time (hours)
S5
129.0588 Da
122.0359 Da
2650
1325
0
Intensity (Counts)
[M-H-CH3CH2OH]⁻
50
100
150
200
m/z (Da)
T
C17H14O7
T1
O
OH
HO
O
O
OH
O
metabolite T
T2
100
Relative Intensity (%)
0
9.2
20
RT (minutes)
T3
C17
329.0661 Da
346.1228 Da
1x10^6
5x10^5
0
Intensity (Counts)
330
340
350
m/z (Da)
T4
1.5x10^7
7.5x10^6
0
Intensity (Counts)
0
24
48
72
96
Incubation Time (hours)
T5
270000
135000
0
Intensity (Counts)
[M-H-CH2O]⁻
[M-H-·CH3]⁻
270
300
330
350
m/z (Da)

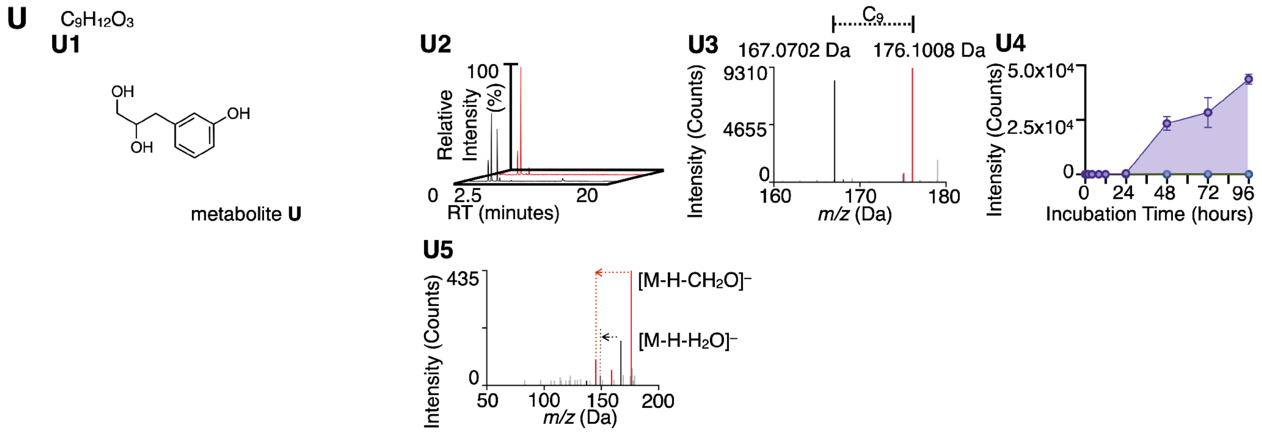


**Figure S4: Summary of metabolomic data for a network of 21 metabolites that result when human gut microbiota deconstruct lignin. (A–U) Panels 1** Structure and molecular formula. **Panels 2** Overlaid extracted ion chromatograms for the $^{12}$C and $^{13}$C *m/z* values. **Panels 3** Overlaid mass spectra for the $^{12}$C and $^{13}$C isotopologs. **Panels 4** Overlaid metabolic profile plots for the $^{13}$C isotopolog across all conditions ($GAM_{+fecal/+lignin}$, $GAM_{-fecal/+lignin}$, $GAM_{+fecal/-lignin}$), depicting average signal intensity at each incubation timepoint (mean ± standard error of the mean, n = 3). For metabolites **C**, **J**, and **O**, metabolic profiles were not obtained for all culture conditions and across all timepoints, which has been noted accordingly. **Panels 5** Overlaid fragmentation mass spectra for the $^{12}$C and $^{13}$C isotopologs, depicting the fragments that correspond to particular metabolites or particular neutral-loss species. Metabolites **D**, **E**, **F**, **K**, **L**, **M**, **N**, **Q**, and **R** show fragmentation consistent with metabolite **T** (Tricin), metabolite **E** shows fragmentation consistent with metabolite **K,** and metabolite **G** shows fragmentation consistent with Metabolite **P**; each of these fragments are denoted in the respective fragmentation spectra. **Panels 6** Metabolite **D** only: Zoomed in overlaid fragmentation mass spectra for the $^{12}$C and $^{13}$C isotopologs of metabolite **D** depicting the fragments that correspond to metabolite **L** and to the lignin subunit 3-(4-hydroxy-3-methoxyphenyl)propan-1,3-diol (1,3VG).

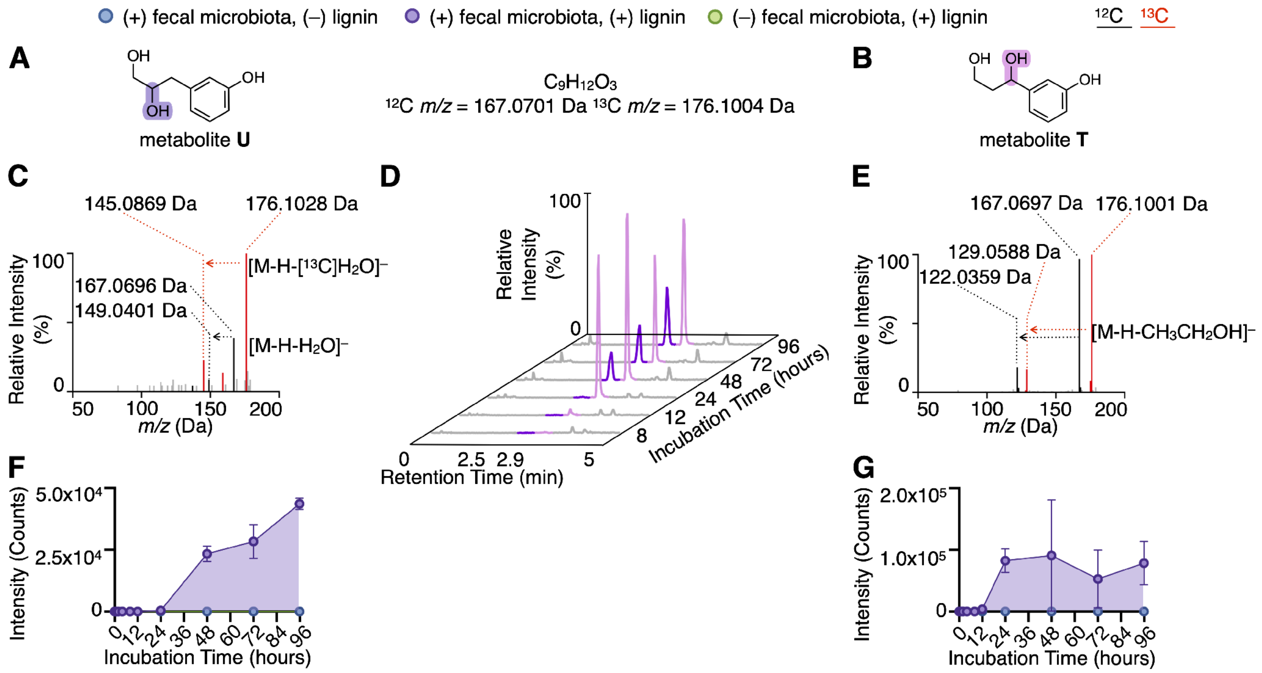


**Figure S5: Different products result from distinct pathways by which human gut microbiota depolymerize lignin. (A)** Structure of metabolite **U**, which results from the hydrolytic depolymerization of metabolite **R (Figure 5R)**. **(B)** Structure of metabolite **S**, which results from the reductive depolymerization of metabolite **Q (Figure 5Q)**. **(C)** Overlaid fragmentation mass spectra of the $^{12}C$ and $^{13}C$ isotopologs of metabolite **U**, where the $^{13}C$ variant was characterized by the neutral-loss of a formaldehyde fragment (176.1028 Da → 145.0869 Da, $[M\text{-}H\text{-}[^{13}C]H_2O]^-$), representative of a 1,2-diol. **(D)** Overlaid extracted ion chromatograms for *m/z* = 176.10 Da, averaged across n = 3 samples for each incubation time. The peaks for metabolite **U** (purple, elution ca. 2.5 minutes) and the peaks for metabolite **S** (pink, elution ca. 2.9 minutes) are characterized by different elution profiles across the incubation period, supporting the hypothesis that they are distinct metabolites that result from different metabolic processes. **(E)** Overlaid fragmentation mass spectra of the $^{12}C$ and $^{13}C$ isotopologs of metabolite **S**, characterized by the neutral-loss of ethanol (176.1001 Da → 129.0588 Da, $[M\text{-}H\text{-}[^{13}C]H_3[^{13}C]H_2OH]^-$; 167.0697 Da → 122.0359 Da, $[M\text{-}H\text{-}CH_3CH_2OH]^-$). **(F)** Metabolic profile of the

$^{13}C$ isotopolog of metabolite **U**, depicting a Production profile. **(G)** Metabolic profile of the $^{13}C$ isotopolog of metabolite **S**, depicting a Production profile.
